# Carbon source diversity drives competitive shifts in multispecies bacterial communities despite attenuated growth enhancement

**DOI:** 10.64898/2026.11.13.766413

**Authors:** Hiroki Ono, Saburo Tsuru, Chikara Furusawa

**Affiliations:** Department of Biological Sciences, Graduate School of Science, The University of Tokyo, 7-3-1 Hongo, Bunkyo-ku, Tokyo 113-0033, Japan; Universal Biology Institute, Graduate School of Science, The University of Tokyo, 7-3-1 Hongo, Bunkyo-ku, Tokyo 113-0033, Japan; Center for Biosystems Dynamics Research, RIKEN, 6-7-1 Minatojima-minamimachi, Chuo-ku, Kobe 650-0047, Japan

**Keywords:** carbon source diversity, interspecies interactions, synthetic microbial communities, community complexity, microbial competition, bacterial growth, resource competition

## Abstract

Bacterial interspecies interactions are often shaped by the growth environment through processes such as resource competition and metabolite exchange. Clarifying the relationship between these interactions and the growth environment is therefore essential to understanding them. Previously, we analysed 28 pairwise coculture systems comprising eight bacterial species across multiple carbon-source environments and found that competitive interspecies interactions, in which the species mutually inhibited growth, became more frequent as carbon source diversity increased. In monospecies cultures, mixtures of carbon sources could also support greater growth than the average observed when the constituent carbon sources were supplied individually, suggesting a benefit beyond increasing the likelihood of providing a usable carbon source. We proposed that this growth enhancement might contribute to the shift towards competitive interactions. However, whether these patterns extend to communities comprising more species remained unclear. Here, we addressed this question in a community comprising the same eight bacterial species. The carbon-source dependence of interactions in the eight-species community was consistent with the pattern observed in pairwise cocultures. In contrast, the growth enhancement associated with carbon-source diversification in monospecies cultures was attenuated in the multispecies community. The persistence of competitive shifts despite attenuated growth enhancement suggests that carbon source diversity acts as a robust driver of competitive interactions across scales of community complexity, offering a basis for relating environmental parameters to interaction patterns in complex microbial communities.

## Introduction

Microorganisms in nature commonly form communities in which multiple species coexist [1]. These communities perform diverse biological and ecological functions, including host digestion [2, 3], metabolism [4], and biogeochemical cycling in ecosystems [5]. Their functions reflect not simply the sum of the traits of individual constituent species, but also interactions among them [6, 7]. Understanding, controlling, and applying microbial community functions therefore requires knowledge of both the traits of constituent species and the interactions among them [8–10].

Understanding bacterial interspecies interactions requires consideration of their carbon-source dependence. Bacteria modify their growth environments through processes including resource uptake and the release of metabolites and other compounds [11–15]. These environmental modifications give rise to diverse interspecies interactions, including growth inhibition through resource competition [16–19] and growth promotion through metabolite provision [20, 21]. Accordingly, bacterial interactions depend on both the species combination and the growth environment [22– 24]. Understanding these interactions therefore requires consideration of the environments in which bacteria grow as well as the species involved.

Despite the strong influence of the growth environment, the environmental conditions that favour bacterial synergism or competition remain poorly understood. In a previous study, we systematically examined the relationship between interspecies interactions and the growth environment in 28 pairwise coculture systems comprising eight bacterial species across 32 environments [24]. Single-carbon-source environments supported diverse interactions, including mutualism, which facilitated the growth of both species, and competition, which inhibited the growth of both species. As the number of carbon sources increased, interactions shifted towards competitive interactions, including both competition and amensalism. We also identified two growth-related effects of increasing carbon source diversity. First, the proportion of bacterial species capable of growth in a given environment increased with the number of carbon sources, consistent with a greater likelihood of providing usable carbon sources. Second, carbon-source mixing could enhance the growth of individual bacterial species: monospecies cultures (mono-cultures) in multi-carbon-source environments frequently reached higher cell densities than the mean observed when the constituent carbon sources were supplied individually. Whereas the first effect concerns whether species can grow, the second concerns the extent of their growth relative to the corresponding single-carbon-source environments. Competitive interactions were also frequently observed in environments in which both species could grow sufficiently. Together, these observations suggested that changes in bacterial growth associated with carbon-source diversification might contribute to the shift towards competitive interactions.

However, how the environmentally dependent interaction patterns identified in pairwise cocultures extend to more complex multispecies communities remains unclear. In multispecies communities, resource consumption and metabolic activity by each constituent species modify the environment and thereby influence multiple community members [25–27]. The diversity of secreted metabolites is also expected to increase with the number of species [28, 29]. This diversification could intensify resource competition [24, 30] or increase opportunities for synergism through cross-feeding [12, 31, 32]. The shift towards competitive interactions with increasing carbon source diversity observed in pairwise cocultures might therefore be strengthened, attenuated, or otherwise altered in multispecies communities. A separate question is whether the growth enhancement associated with carbon-source mixing in mono-cultures persists in multispecies communities. The metabolic activities of multiple species may alter growth responses to carbon-source mixing, potentially changing the extent of this enhancement in multispecies communities. Determining whether knowledge from simple systems can inform our understanding of more complex multispecies communities therefore requires direct tests of the extent to which these patterns persist.

Here, we tested whether the shift towards competitive interactions observed in pairwise cocultures and the growth enhancement observed in mono-cultures with increasing carbon source diversity also occur in multispecies bacterial communities. Using the same eight bacterial species as in the previous study, we cultured a community containing all eight species (full-culture), eight communities each lacking a single species (dropout-cultures), and mono-cultures of each species across 19 carbon-source environments (**Fig. 1A, B**). We quantified interactions in the multispecies community by treating each species as the focal species and evaluating the net effects between it and the resident community comprising the other seven species (**Fig. 1C, D**). The shift towards competitive interactions with an increasing number of carbon sources observed in pairwise cocultures was also evident in multispecies communities. Competitive species–community interactions were more likely in environments in which both the focal species and the resident community could grow sufficiently. In contrast, although mono-cultures showed growth enhancement associated with carbon-source mixing, as in the previous study, this enhancement was attenuated in dropout-cultures and full-cultures. These results reveal a dissociation between the two phenomena: community complexity buffers the growth enhancement driven by carbon-source mixing while leaving the shift towards competitive interactions largely intact, suggesting that carbon source diversity acts as a robust driver of competitive interactions across scales of community complexity.

**Figure 1.**
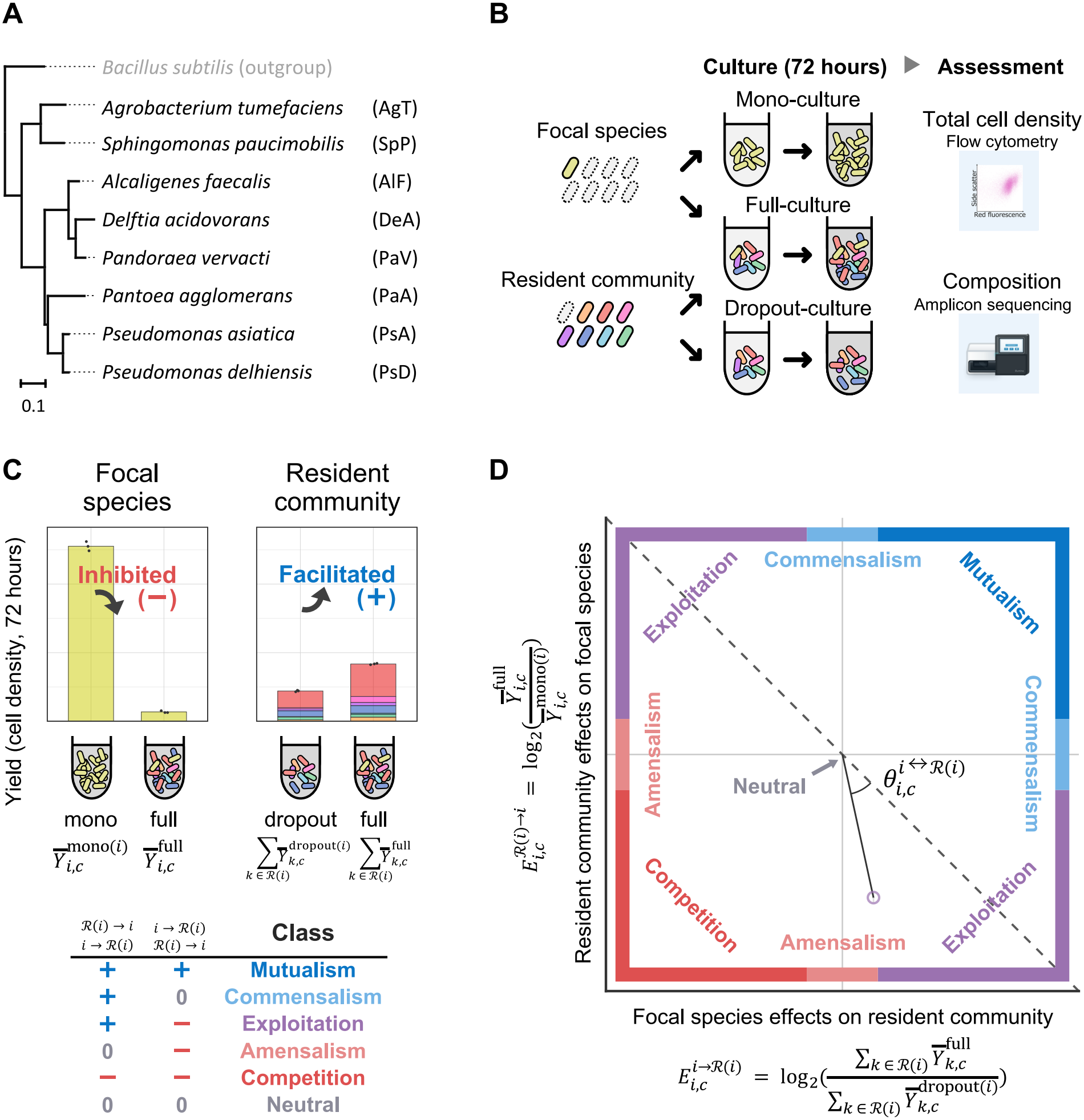
Experimental design and effect-plane framework for full-culture and dropout-culture communities. Schematic overview of the experimental design and species–community interaction framework. (**A**) Phylogenetic relationships among the eight bacterial species analysed, with *Bacillus subtilis* as the outgroup. The 16S rRNA gene phylogeny was reused from the previous study [24]. Species abbreviations used throughout the manuscript are shown in parentheses on the right, and the scale bar is shown at the lower left. (**B**) Experimental design for investigating interspecies interactions. In each carbon-source environment, we prepared mono-cultures of individual species, a full-culture containing all eight species, and dropout-cultures each lacking one species, then measured cell density and species composition at 72 hours. (**C, D**) Evaluation of the two directional net effects that define species–community interactions. These are the change in focal-species yield in full-culture relative to mono-culture, 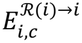 (effect of the resident community on the focal species; y-axis in **D**), and the change in resident-community yield in full-culture relative to the dropout-culture lacking the focal species, 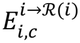 (effect of the focal species on the resident community; x-axis in **D**). An interaction class was assigned from the signs of these two effects (+, 0, or −). The effects 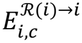 and 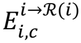were plotted on the vertical and horizontal axes, respectively. The angle between the line segment connecting a plotted point to the origin and the line *y* = −*x* defines interaction type, 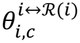, whereas the distance from the origin defines interaction strength, 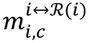. Error bars represent the standard deviation across three biological replicates.

## Materials and Methods

### Bacterial strains

We used eight non-fluorescently labelled bacterial strains from the same species set used in the previous study of interspecies interactions in pairwise cocultures [24] (**Fig. 1A**). Higher-level taxonomy, 16S rRNA gene copy numbers, reference sequences for species assignment, and strain names are provided in **Supplementary Table S1**.

### Culture medium

M9-based media were prepared immediately before use to control the number of carbon sources in the growth environment. The following components were combined, and the final volume was adjusted to 1.0 L with pure water: 100 mL of M9 salts solution containing NaCl (5.00 mg/mL), NH_4_Cl (10.00 mg/mL), KH_2_PO_4_ (30.00 mg/mL), and Na_2_HPO_4_ (67.78 mg/mL), adjusted to pH 7.2 with 8.0 M NaOH; 1.0 mL of trace-elements solution containing H_3_BO_3_ (24.8 μg/mL), MnCl_2_ (16.0 μg/mL), CoCl_2_ (7.2 μg/mL), (NH_4_)_6_Mo_7_O_24_ (3.4 μg/mL), ZnSO_4_ (2.8 μg/mL), and CuSO_4_ (2.4 μg/mL); 1.0 mL of MgSO_4_ solution (240.70 mg/mL); 1.0 mL of CaCl_2_ solution (11.10 mg/mL); and 1.0 mL of FeSO_4_ solution (1.52 mg/mL). We examined environments containing one, two, four, eight, or 16 carbon source types, supplied individually or in hierarchical combinations at a total concentration of 1.0 mg/mL. We selected 19 of the 32 carbon-source environments used in the previous study of pairwise cocultures [24]; their compositions are shown in **Table 1**. The previous study confirmed that a total carbon-source concentration of 1.0 mg/mL limited mono-culture growth in many species–environment combinations. A further 18 environments were included in the supplementary analysis described below (**Supplementary Table S2**). Information on all suppliers is provided in **Supplementary Table S3**. Unless otherwise noted, all solutions were prepared in pure water (**Supplementary Methods S1**).

**Table 1.** Carbon-source environments for investigating interspecies interactions.

| **Name of environment** | **Component carbon sources** | **No. of carbon sources** |
|---|---|---|
| Cellobiose | Cellobiose | 1 |
| Sorbitol | Sorbitol | 1 |
| Serine | Serine | 1 |
| Raffinose | Raffinose | 1 |
| Uridine | Uridine | 1 |
| Glycerol | Glycerol | 1 |
| Citrate | Citrate | 1 |
| Proline | Proline | 1 |
| 2-1 | Cellobiose + sorbitol | 2 |
| 2-2 | Serine + raffinose | 2 |
| 2-3 | Uridine + glycerol | 2 |
| 2-4 | Citrate + proline | 2 |
| 4-1 | 2-1 + 2-2 | 4 |
| 4-2 | 2-3 + 2-4 | 4 |
| 4-3 | Alanine + glutamine + mannitol + isoleucine | 4 |
| 4-4 | Glucose + ribose + acetate + succinate | 4 |
| 8-1 | 4-1 + 4-2 | 8 |
| 8-2 | 4-3 + 4-4 | 8 |
| 16-1 | 8-1 + 8-2 | 16 |

### Preculture, community assembly, and endpoint sampling

Each species was precultured by inoculating M9-based liquid medium containing 0.5 mg/mL glucose and 0.5 mg/mL succinate with cells from a frozen stock (200 μL, 96-well plate, 33 °C, 500 rpm, 48 hours), using plate shakers (MBR-034P, TAITEC, Saitama, Japan). Cultures were then diluted on the basis of OD_595_ (optical density at 595 nm) measured by a plate reader (Infinite F200, Tecan, Männedorf, Switzerland) and the known growth rate of each strain, followed by a further 18 hours of preculture under the same conditions. After preculture, the cell density of each strain was measured by flow cytometry (FACS Aria III Cell Sorter, BD, New Jersey, USA), and inoculation suspensions were adjusted to a common cell density. In each carbon-source environment, we prepared a full-culture containing all eight species, eight dropout-cultures each lacking one species, and mono-cultures of each of the eight species. The total initial inoculum was standardised to 20,000 cells/well for full-culture, dropout-culture, and mono-culture, irrespective of species number, and cultures were incubated for 72 hours (200 μL, 96-well plate, 33 °C, 500 rpm). Cultures collected at 72 hours were used to measure total cell density by flow cytometry and for 16S rRNA gene amplicon sequencing. To assess growth near the endpoint, we measured OD_595_ at 66, 69, and 72 hours and evaluated values corrected against blank wells (**Supplementary Fig. S1**). Samples were flagged when OD_595_ changed by more than 10% between adjacent measurements, using the higher OD_595_ value in each interval as the reference. Cell density, composition, and representative yield panels for each environment and culture type are provided in **Supplementary Fig. S2** and **Supplementary Table S4**.

### 16S rRNA gene amplicon library preparation

Templates for 16S rRNA gene amplicon sequencing were prepared from aliquots of endpoint cultures using a method based on a previous study [33]. We added 3 μL of lysozyme solution (50 mg/mL lysozyme, 20 mM Tris-HCl, pH 8.0) to 15 μL of culture from each sample and incubated the mixture at 37 °C for 2 hours. We then added 3 μL of proteinase K solution (0.70 mg/mL recombinant proteinase K, 120 mM Tris-HCl, pH 8.0, 0.105 mM CaCl_2_, and 1.75% (v/v) glycerol), incubated the mixture at 55 °C for 3 hours, and heated it at 95 °C for 10 minutes. The mixture was then vortexed for 10 minutes, and the resulting lysate was used as the PCR template. The V4 region of the prokaryotic 16S rRNA gene was amplified by first-round PCR using primers modified from 515F/806rB [34, 35] with added Nextera overhang sequences. The primer sequences were as follows: forward primer, 5′-TCGTC GGCAG CGTCA GATGT GTATA AGAGA CAGGT GCCAG CAGCC GCGGT AA-3′; reverse primer, 5′-GTCTC GTGGG CTCGG AGATG TGTAT AAGAG ACAGG GACTA CCAGG GTATC TAAT-3′. The first-round PCR mixture contained 11 μL of 44-fold-diluted lysate, 0.75 μL of 10 μM forward primer, 0.75 μL of 10 μM reverse primer, and 12.5 μL of KOD One Dye-containing master mix. After initial denaturation at 98 °C for 10 seconds, PCR was performed for 40 cycles of 98 °C for 10 seconds, 55 °C for 5 seconds, and 68 °C for 1 second, followed by extension at 68 °C for 10 seconds and holding at 4 °C. AMPure XP beads (20 μL) were added to 25 μL of first-round PCR product, and the products were bound, washed, and dried according to the manufacturer’s recommended procedure. DNA was eluted in 53 μL of 10 mM Tris-HCl, pH 8.0, and 50 μL of the eluate was recovered. Second-round PCR (indexing PCR) was performed using Nextera XT index primers. The PCR mixture contained 2.5 μL of purified first-round PCR product, 2.5 μL of Nextera XT index 1 primer, 2.5 μL of Nextera XT index 2 primer, 12.5 μL of 2× Gflex PCR Buffer, 0.5 μL of Tks Gflex DNA polymerase, and 4.5 μL of Milli-Q water. After initial denaturation at 94 °C for 1 minute, PCR was performed for 12 cycles of 98 °C for 10 seconds, 55 °C for 15 seconds, and 68 °C for 15 seconds, followed by holding at 4 °C. AMPure XP beads (28 μL) were then added to 25 μL of second-round PCR product, and purification was performed according to the manufacturer’s recommended procedure. DNA was eluted in 53 μL of 10 mM Tris-HCl, pH 8.0, and 50 μL of the eluate was recovered. Purified second-round PCR products were quantified using a Quant-iT dsDNA BR Assay Kit and a Tecan plate reader. Each sample library was diluted to 4 nM with 10 mM Tris-HCl, pH 8.0, and the final library pool was prepared by combining 5 μL of each diluted library.

### MiSeq sequencing and amplicon processing

The prepared libraries were sequenced on an Illumina MiSeq (MiSeq System, Illumina, Inc., San Diego, CA, USA) using a MiSeq Reagent Kit v3 (150 Cycles; 25M), and raw FASTQ data were processed using DADA2 [36]. Reads were filtered using a truncation length of 70 bp for both forward and reverse reads, a maximum of zero ambiguous nucleotides, maximum expected errors of 2 for both reads, a truncation quality score of 2, and PhiX read removal. Error models were learned separately for forward and reverse reads in each run, and amplicon sequence variants (ASVs) were inferred after dereplication. In mergePairs, justConcatenate was set to TRUE, yielding 150-bp ASV sequences comprising a 70-bp forward read, a 10-N spacer, and a 70-bp reverse read. Sequence tables from each run were combined, and chimeras were removed using the consensus method. Run-specific quality-profile evaluations, independently learned DADA2 error models, read tracking, and read-depth balance after chimera removal are summarised in **Supplementary Table S5**. Taxonomic assignment used a custom reference FASTA generated from reference sequences for the eight target species, and ASVs were aggregated to the species level on the basis of species_code. Unassigned ASVs were retained for quality control but excluded from the denominator for species-level relative abundance. Species assignment, the unassigned-read fraction, and the handling of dropout-contamination ranking are described in **Supplementary Table S6**.

### Flow-cytometric total cell-density measurement

Total cell density was measured by flow cytometry using propidium iodide (PI)-stained cultures and Fluoresbrite Polychromatic Red Microspheres of known concentration (1.0 μm; PC Red). Cultures were adjusted with phosphate-buffered saline (PBS) to final concentrations of 50% ethanol and 2.5 μg/mL PI, then incubated at room temperature for 30 minutes before measurement. Flow cytometry was performed using a FACS Aria III Cell Sorter, with event data acquired for each sample at a flow rate of 2.0 for 1 minute. PI-positive cell populations, PC Red, and debris were identified on the basis of side scatter intensity, red fluorescence intensity, and green fluorescence intensity. Total cell density was calculated from the ratio of PI-positive-cell events to PC Red events, the known number of PC Red beads added, and the culture aliquot volume. Fluorescence intensity gates and the relationship between OD_595_ and cell density are shown in **Supplementary Fig. S3** and **Supplementary Fig. S4**, respectively.

### Abundance estimation

Species-specific relative abundance was calculated from species-level read counts for each sample. To correct for biases arising from differences in 16S rRNA gene copy number among species, each species’ relative abundance was divided by its corresponding copy number. The corrected values were then renormalised to sum to one within each sample. Species-specific absolute abundance was estimated by multiplying this copy-number-corrected relative abundance by the total cell density measured by flow cytometry in the same sample [37–39].

### Species–community interaction

We defined species–community interactions as the bidirectional effects between a focal species *i* and the resident community. The set of all eight species was denoted by *S*, and the resident community excluding focal species *i* by & *c*ℛ (*i*) = *S \* {*i*}. For each focal species *i* and environment *c*, we quantified the effect of the resident community ℛ (*i*) on *i* and the effect of *i* on the total yield of the resident community ℛ(*i*) as follows.

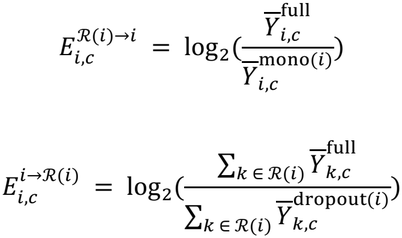

Here, 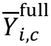 and 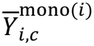 represent the mean yields of species *i* at 72 hours across three biological replicates in full-culture and mono-culture, respectively. 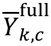 and 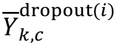 denote the mean cell densities of resident species *k* across three biological replicates in full-culture and the corresponding dropout-culture, respectively. In both species– community and species–species interaction analyses, yields were analysed as absolute cell densities on the original scale. For each directional effect, replicate yields were compared using a two-sided Welch two-sample *t*-test [40]. Within each analysis scale and interaction analysis, *P* values from both directions across the included environments were combined and adjusted using the Benjamini–Hochberg method [41]; the adjusted *P* values are denoted as *q* values. A directional effect was classified as positive when *q* < 0.05 and the effect was positive (+), as negative when q < 0.05 and the effect was negative (−), and as neutral otherwise (0). The ordered sign combination, with the effect of the focal species on the resident community first and the reverse effect second, defined six interaction classes: mutualism (+, +), commensalism (+,0) or (0, +), exploitation (+, −) or (−, +), amensalism (−,0) or (0, −), competition (−, −), and neutral (0,0) [8, 42, 43].

Following previous studies [22, 24], we quantified interaction type as a measure of synergism based on the two directional effects, as follows.

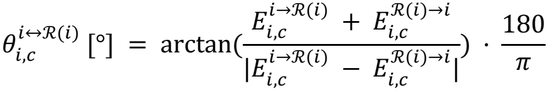

This metric is the angle between the line segment connecting a plotted point to the origin in **Fig. 2A** and the line *y* = −*x*. When the denominator is zero because the two effects are equal, the metric is defined as 90°, −90°, or 0°, according to their sign. Larger values indicate more synergistic interactions, whereas smaller values indicate more competitive interactions.

**Figure 2.**
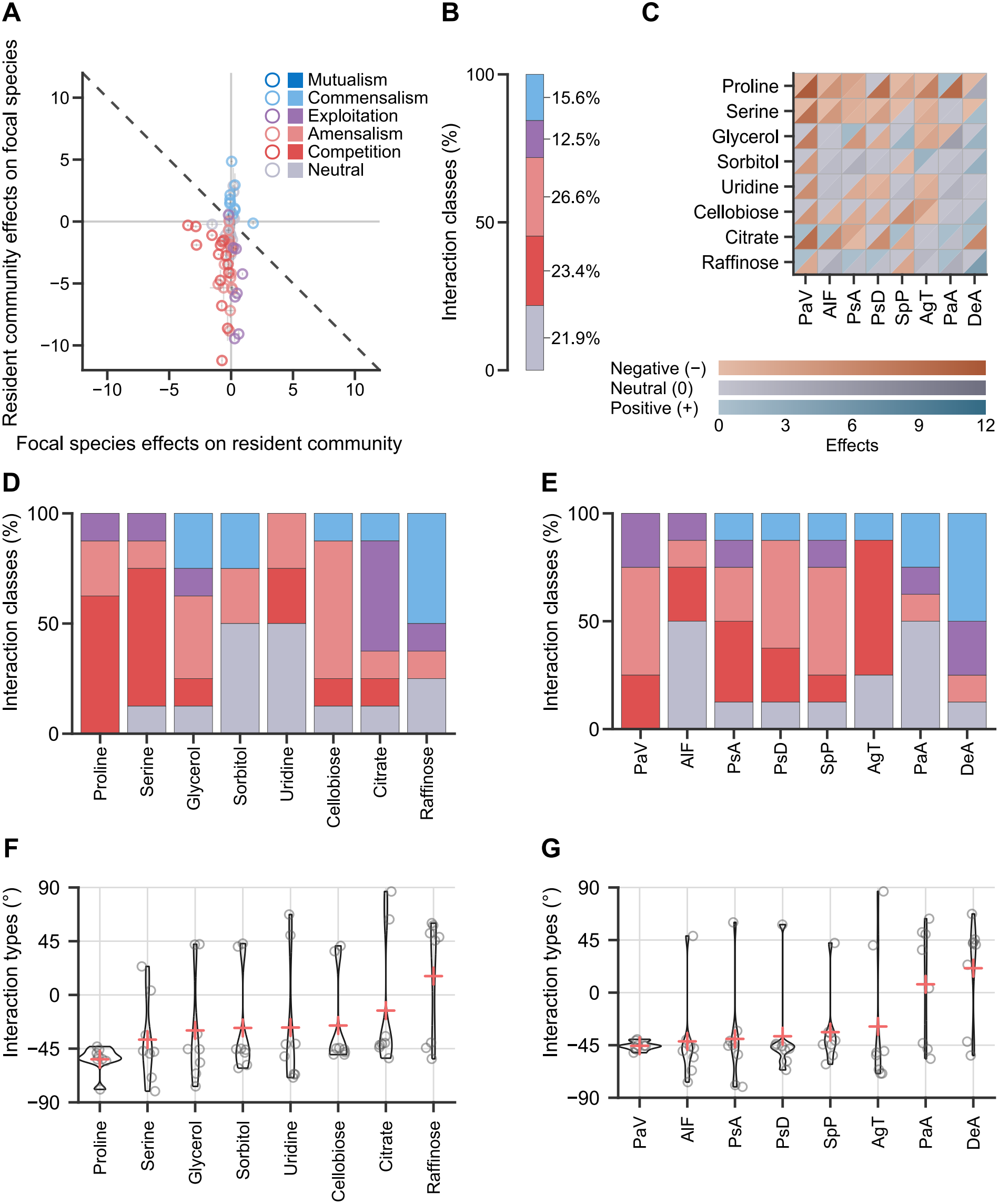
Species–community interactions in single-carbon-source environments. Species–community interaction patterns across 64 combinations of eight single-carbon-source environments and eight focal species. (**A**) Effect plane showing the effect of the focal species on the resident community, 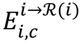 (x-axis; log_2_ ratio between resident-community yield in full-culture and in the dropout-culture lacking the focal species), and the effect of the resident community on the focal species, 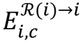 (y-axis; log_2_ ratio between focal-species yield in full-culture and mono-culture). Each point represents one focal species–environment combination, with colour indicating the interaction class assigned from the signs of the two directional effects. Error bars represent the standard deviation across three biological replicates. (**B**) Fractions of interaction classes across the same 64 combinations shown in **A**. (**C**) Split-triangle heatmap of directional effects for the eight environments and eight focal species. The upper-left and lower-right triangles of each cell indicate the effects of the focal species on the resident community and of the resident community on the focal species, respectively. Blue, red, and grey indicate positive, negative, and neutral effects, respectively; colour intensity indicates the magnitude of the corresponding directional log_2_ effect. (**D, E**) Interaction-class fractions by carbon-source environment (**D**) and focal species (**E**), ordered by their respective mean interaction types, 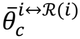 and 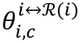. (**F, G**) Jittered violin plots showing the distribution of interaction types, 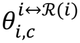, by carbon-source environment (**F**) and focal species (**G**). Circular points represent individual interaction types, 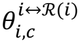, and cross-shaped points indicate the respective mean interaction types, 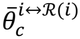 and 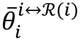, for each carbon source or focal species. Bacterial-species abbreviations are provided in **Fig. 1A**.

Following a previous study [22], we also defined interaction strength using the following equation.

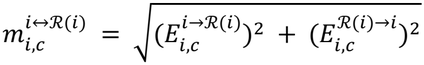

This metric corresponds to the Euclidean distance between a plotted point and the origin in **Fig. 2A**.

### Species–species interaction

Species–species interactions represent species-level directional effects within the same multispecies community. In growth environment *c*, we defined the effect of target species *j* on focal species *i* and the effect of focal species *i* on target species *j* as follows.

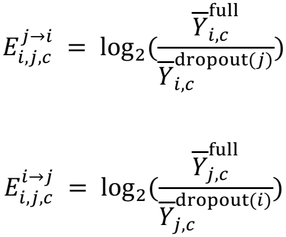

Both directional effects were evaluated using the procedures described above and classified as +, −, or 0 to assign an interaction class. Interaction type and interaction strength were calculated as follows.

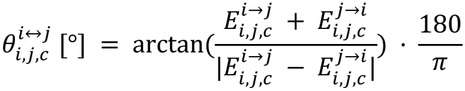

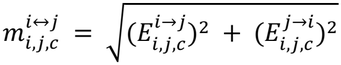

### Association between endpoint cell density and competitive interactions

To examine the association between endpoint cell density and competitive interactions, we used the mean cell density at 72 hours of the focal species in mono-culture and the corresponding resident community in dropout-culture. The analysis included eight focal species in each of 19 carbon-source environments, yielding 152 species–community interactions. Each cell-density measure was divided into Low and High groups at its median across the 152 interactions, defining four cell-density quadrants. Interactions assigned to the Competition or Amensalism class were classified as competitive; those assigned to Mutualism, Commensalism, Exploitation, or Neutral were classified as non-competitive.

We tested the global association between cell-density quadrant and competitive-interaction status using the Pearson chi-square statistic for the 4 × 2 table, with *P* values calculated from 10,000 environment-block permutations. In each permutation, the complete ordered vector of eight interaction outcomes from one environment was reassigned to another while retaining focal-species positions. This procedure preserves the dependence among focal-species observations within each environment and the outcome frequencies specific to each focal species. We then used one-sided environment-block permutation tests to assess whether the competitive-interaction fraction in each quadrant exceeded the overall fraction.

### Analysis of growth enhancement associated with carbon-source mixing

We evaluated growth enhancement associated with carbon-source mixing using cell density or OD_595_ as the yield measure. Following the previous study [24], we compared Group 1, representing growth in a multi-carbon-source environment, with Group 2, representing the mean growth when the constituent carbon sources were supplied individually.

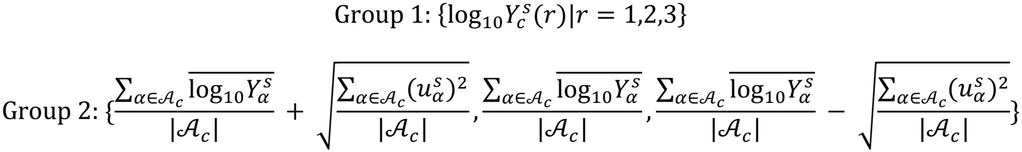

Here, 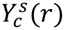 represents the yield of culture unit *s* after 72 hours in a multi-carbon-source environment *c* for biological replicate *r*. The set of single carbon sources comprising environment *c* is denoted by *A*_*c*_, and the number of carbon sources by |*A*_*c*_|. Culture unit *s* represents one mono-culture of a focal species (eight types), one dropout-culture (eight types), or the full-culture (one type). For each constituent single carbon source 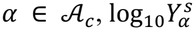 represents the mean log_10_ yield across three replicates, and 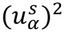 represents the unbiased variance of log_10_ yield across those replicates. Thus, Group 1 comprises three observed replicate values of log_10_ yield in the multi-carbon-source environment. Group 2 is a theoretical group comprising three log_10_ yield values, with a mean corresponding to the mean growth when the constituent carbon sources are supplied individually and a variance reflecting variation among those replicates. We used a two-sided *t*-test [40] to test for a difference between the group means. The resulting *P* values were adjusted using the Benjamini–Hochberg method [41] to control the false discovery rate. Results were classified as High when *q* < 0.05 and the observed mean exceeded the expected mean, as Low when q < 0.05 and the observed mean was lower, and as Same otherwise. For OD_595_, we analysed the 18 environments in **Supplementary Table S2** in addition to the 19 environments in which cell density was measured. The observed and expected groups in the OD_595_ analysis were constructed using untransformed, blank-corrected values.

In single-carbon-source environments, we compared community yields with predictions from constituent mono-cultures. For full-culture, the observed and theoretical groups were as follows.

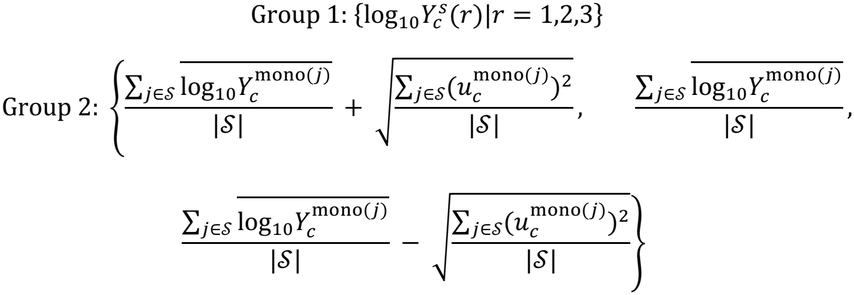

For dropout-culture lacking species *i*, *S* in Group 2 was replaced by ℛ(*i*) . Welch testing and High/Same/Low classification followed the same procedure, with BH correction applied separately by yield measure and culture type.

### Comparison with prior pairwise interactions

Directional effects and interaction classes from Supplementary Table S8 of Ono et al. [24] were matched to the current species–species estimates by environment and species pair (19 environments × 28 pairs). Differences in interaction strength, calculated as current minus prior estimates, were analysed using a linear mixed model fitted by maximum likelihood, with crossed random intercepts for environment and species pair. The overall mean difference was tested against zero by a likelihood-ratio comparison with a model omitting the fixed intercept, and its 95% confidence interval was obtained by likelihood profiling. Agreement in interaction class and agreement in the two directional signs were summarised separately (**Supplementary Table S9**).

### Computational environment

Analyses were performed in R [44] and Python [45], with NumPy [46], pandas [47], and SciPy [48] used for supporting data processing and calculations. DADA2 [36], phyloseq [49], Biostrings [50], and pwalign [51] were used for 16S amplicon processing, whereas base R, tidyverse [52], vegan [53], and ggplot2 [54] were used for statistical analyses and visualisation.

## Results

### Species–community interactions vary across single-carbon-source environments

Across 19 carbon-source environments, we cultured the same eight bacterial species used in the previous study [24] as a full-culture containing all eight species, eight dropout-cultures each lacking one species, and mono-cultures of each species (**Fig. 1A, B**). Irrespective of the number of carbon sources, the total carbon-source concentration was standardised to 1 mg/mL, a concentration previously confirmed to limit mono-culture growth in many species– environment combinations. To evaluate interspecies interactions in the eight-species community, we defined species– community interactions as the effects between a focal species and the resident community comprising the other seven species (**Materials and Methods**). In growth environment *c*, the effect of the resident community ℛ(*i*) on focal species *i*, denoted by 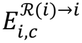, was calculated as the log_2_ ratio of full-culture to mono-culture yield, using the mean yields of the focal species in mono-culture, 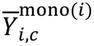, and full-culture, 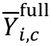 each based on three biological replicates. The effect of focal species *i* on the resident community ℛ(*i*), denoted by 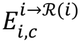, was calculated as the log_2_ ratio of full-culture to dropout-culture yield, using the mean total yields of the resident community in the dropout-culture lacking the focal species, 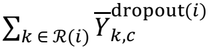, and full-culture, 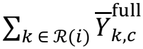, each based on three biological replicates (**Fig. 1C, D**). Unlike effects measured in pairwise cocultures, these estimates combine direct and indirect effects in the multispecies context.

In single-carbon-source environments, species–community interactions ranged from synergistic to competitive, depending on the carbon-source environment and focal species, as in the previous study of pairwise interspecies interactions [24] (**Fig. 2A–C**). Among the 64 focal species–environment combinations, 15 interactions were classified as competition (23.4%), 17 as amensalism (26.6%), eight as exploitation (12.5%), 10 as commensalism (15.6%), and 14 as neutral (21.9%); mutualism was not detected.

To identify factors associated with variation in interspecies interactions, we summarised the observed interactions by growth environment and bacterial species. Interspecies interactions were strongly influenced by the supplied carbon sources (*P* = 2.0 × 10^−3^, chi-square test with Monte Carlo estimation [55], 10^5^ iterations; **Supplementary Table S7**). To quantify the species–community interaction in growth environment *c* for focal species *i*, we used interaction type, 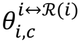, as a measure of synergism. This metric is the angle relative to the dashed line in **Fig. 1D** and is equivalent to the metric used in the previous study of pairwise cocultures (**Materials and Methods**; **Fig. 1D**) [24]. When the two groups (here, the focal species and resident community) exert positive effects on each other, 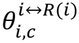 increases to a maximum of 90°; when they exert negative effects on each other, 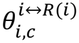 decreases to a minimum of −90°. We ordered single-carbon-source environments by their mean interaction type, 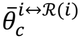. Proline was associated with the most competitive interactions (leftmost column in **Fig. 2D, F**; 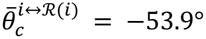), whereas raffinose was associated with the most synergistic interactions (rightmost column in **Fig. 2D, F**; 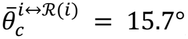).

Interspecies interactions also differed among focal species. When focal species were ordered by mean interaction type, DeA and PaA showed many synergistic interactions (the two rightmost columns, with 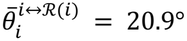 and 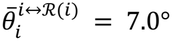, respectively), whereas many other species showed competitive interactions (left columns; for example, PaV, 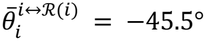; **Fig. 2E, G**). A chi-square test also confirmed a significant association between focal species and interaction class (*P* = 0.05, chi-square test with Monte Carlo estimation [55], 10^5^ iterations; **Supplementary Table S8**). Together, these findings show that both carbon source and species identity influence interspecies interactions in multispecies communities, as observed in pairwise cocultures.

### Carbon source diversity reshapes species–community interactions

We next examined how species–community interactions changed with the number of carbon sources by expanding the analysis to multi-carbon-source environments. The previous study showed that interactions in pairwise cocultures shifted towards competitive interactions as the number of carbon sources increased [24]; we asked whether this tendency also occurs in multispecies communities. As in that study, carbon sources were mixed in equal masses at a total concentration of 1.0 mg/mL, matching that of the single-carbon-source environments.

Across the 19 environments, species–community interactions became more competitive as the number of carbon sources increased (**Fig. 3A**). Competition and amensalism together accounted for 50% of interactions in single-carbon-source environments (leftmost column of the bottom row in **Fig. 3A**) and 100% in the 16-carbon-source environment (rightmost column of the bottom row in **Fig. 3A**). Analysis of interaction type, 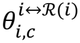, revealed a weak negative correlation with the number of carbon sources (Spearman’s rank correlation: *ρ* = −0.19, *P* = 0.02, **Fig. 3B**). To assess the influence of neutral interactions on this relationship, we repeated the analysis after excluding them; the weak negative correlation remained significant (Spearman’s rank correlation: *ρ* = −0.21, *P* = 0.02, **Supplementary Fig. S5**). To address non-independence among observations, we also averaged interaction types across focal species within each environment and used an environment-level permutation test to evaluate Spearman’s rank correlation *ρ* between the 19 environment means and the number of carbon sources. The relationship based on all interactions in **Fig. 3B** remained negative (Spearman’s rank correlation: *ρ* = −0.64, permutation *P* = 4.3 × 10^−3^; 95% environment-cluster bootstrap confidence interval (CI) [−0.90, −0.22]), as did the relationship after excluding neutral interactions in **Supplementary Fig. S5** (Spearman’s rank correlation: *ρ* = −0.53, permutation *P* = 0.02 ; 95% environment-cluster bootstrap CI [−0.86, −0.07]). These results show that the tendency towards more competitive interactions with increasing carbon source diversity also occurs in multispecies communities.

**Figure 3.**
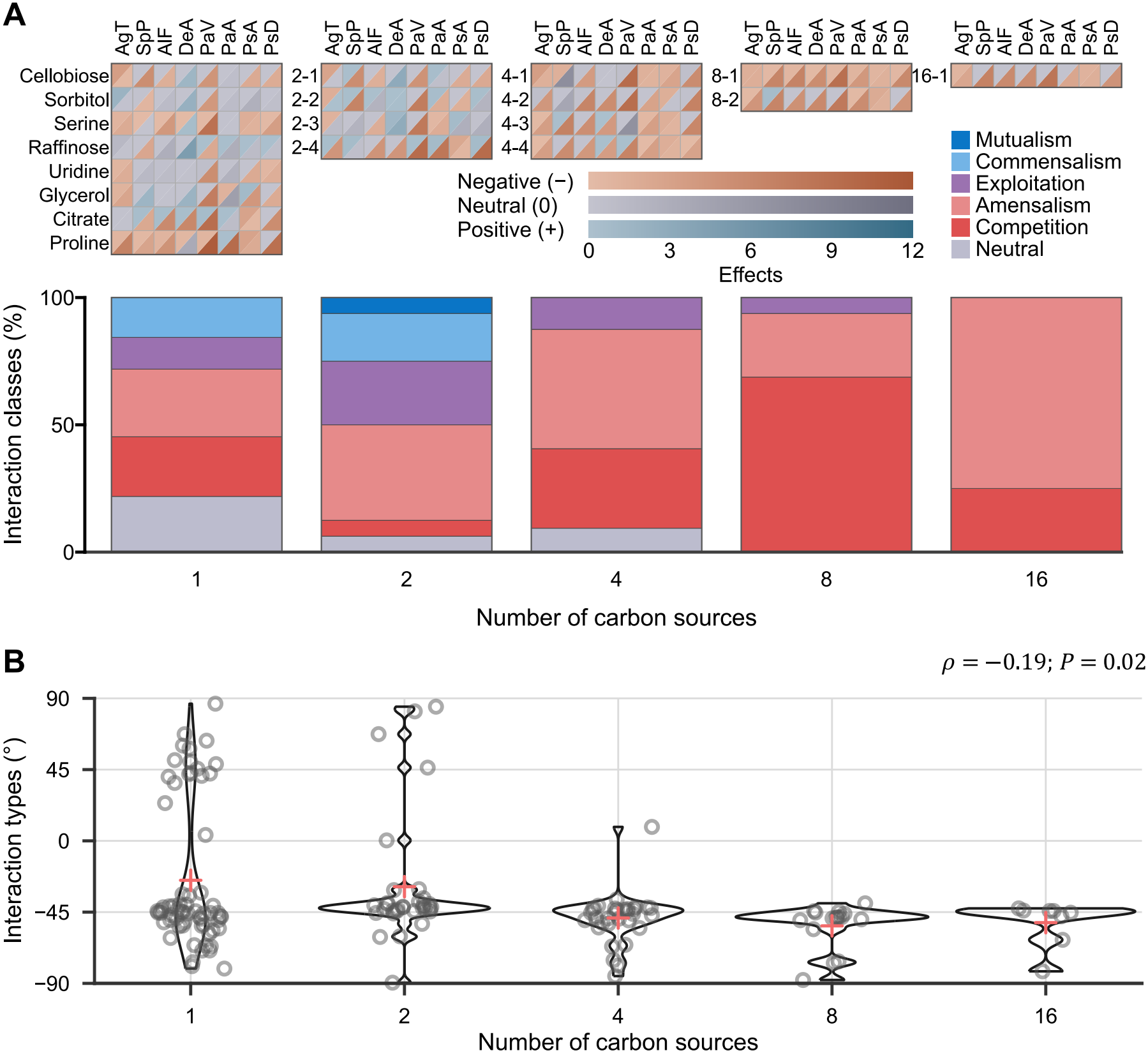
Species–community interactions across carbon source diversity levels. Relationship between species–community interactions and the number of carbon sources across 19 environments, comprising 19 × 8 = 152 focal species–environment combinations. (**A**) Directional effects and interaction-class fractions by number of carbon sources. The upper panel shows split-triangle heatmaps of directional effects for each focal species–environment combination, grouped by number of carbon sources (1, 2, 4, 8, or 16). The upper-left and lower-right triangles of each cell indicate the effects of the focal species on the resident community and of the resident community on the focal species, respectively. Blue, red, and grey indicate positive, negative, and neutral effects, respectively; colour intensity indicates the magnitude of the corresponding directional log_2_ effect. Each heatmap row represents the carbon sources supplied. Bacterial-species and carbon-source abbreviations are provided in **Fig. 1A** and **Table 1**, respectively. The lower panel shows interaction-class fractions for the same combinations, grouped by number of carbon sources. The leftmost plot reproduces the single-carbon-source results in **Fig. 2B, C**. (**B**) Jittered violin plots showing the distribution of interaction types, 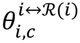, by number of carbon sources. Each point represents one focal species–environment combination, each violin shows the distribution, and cross-shaped points indicate the mean interaction type at each number of carbon sources. Spearman’s rank correlation *ρ* and its *P* value, calculated using 152 points, are shown in the upper-right corner.

### Endpoint growth across carbon-source environments is associated with species–community interactions

The increase in competitive interactions may be associated with growth of the focal species and resident community. In our previous study [24], we identified two potential ways in which increasing carbon source diversity could promote bacterial growth. First, increasing the number of carbon sources could increase the likelihood of providing resources usable by a given species and thus the proportion of species capable of growth. Second, combining carbon sources could enhance the growth of individual species beyond the mean observed when the constituent carbon sources were supplied individually. Competitive interactions also occurred more frequently in environments in which both species could grow sufficiently. We therefore examined endpoint growth in mono-cultures and dropout-cultures across carbon-source environments and its relationship with species–community interactions. Mono-culture endpoint cell density increased with the number of carbon sources (Spearman’s rank correlation: *ρ* = 0.40, *P* = 3.3 × 10^−19^, **Fig. 4A**), whereas no significant trend was observed for dropout-cultures (Spearman’s rank correlation: *ρ* = 0.08, *P* = 0.09, **Fig. 4B**). In a supplementary analysis, we averaged the three replicates within each environment × species condition on the log_10_ scale, then averaged the condition means across the eight species within each environment. We assessed Spearman’s rank correlation *ρ* between the 19 environment means and the number of carbon sources using an environment-level permutation test. This analysis likewise showed a positive relationship for mono-cultures (Spearman’s rank correlation: *ρ* = 0.75, permutation *P* = 2.8 × 10^−4^ ; 95% environment-cluster bootstrap CI [0.47, 0.10]) and no significant relationship for dropout-cultures (Spearman’s rank correlation: *ρ* = 0.20, permutation *P* = 0.42; 95% environment-cluster bootstrap CI [−0.31, 0.61]).

**Figure 4.**
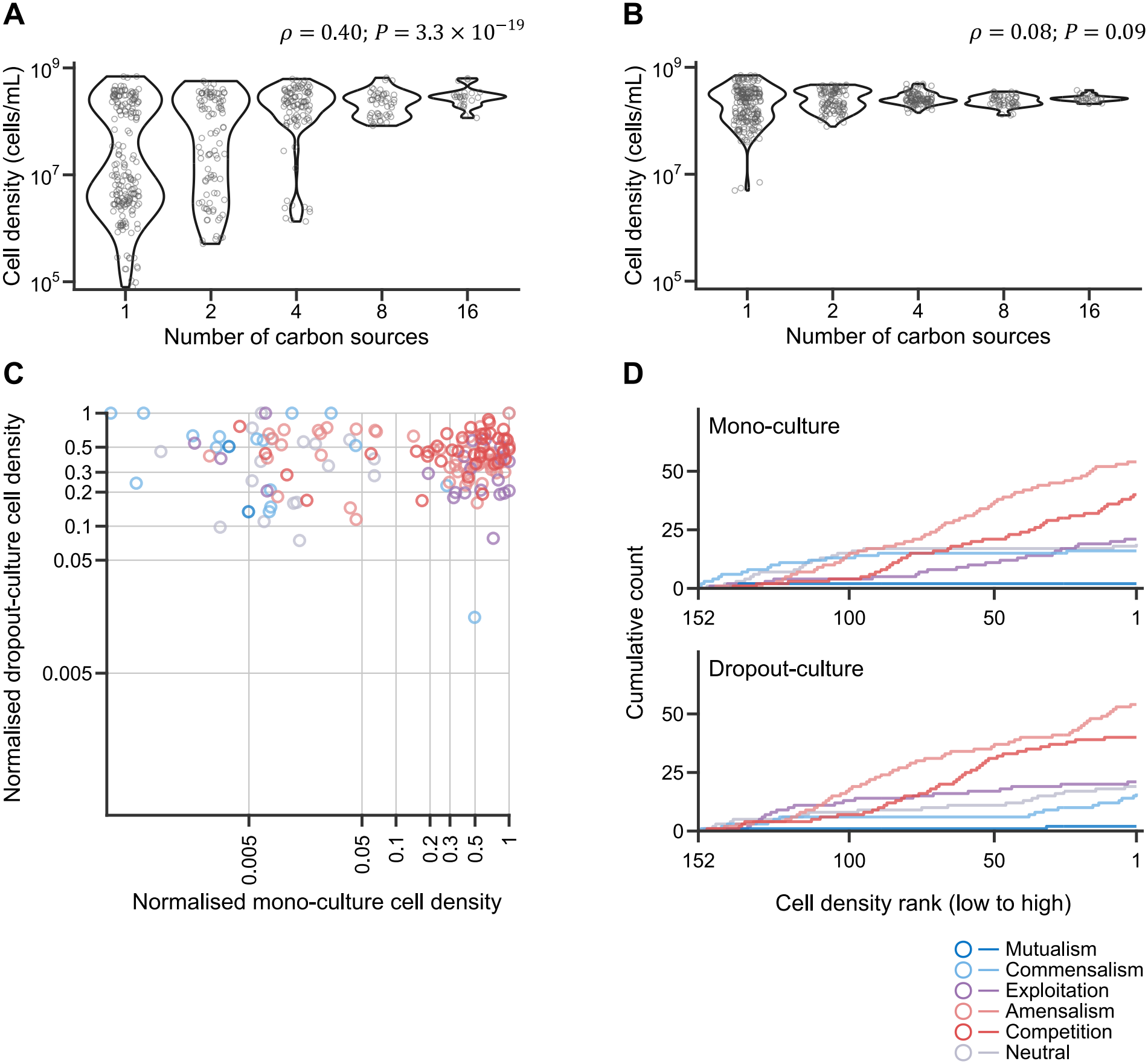
Endpoint growth and species–community interactions across carbon source diversity levels. Relationship between endpoint cell density, carbon source diversity, and species–community interactions. (**A, B**) Cell density at 72 hours by number of carbon sources in mono-cultures (**A**) and dropout-cultures (**B**). Each point represents one sample observation, and each violin shows the distribution; 456 points are plotted on a log_10_ scale. Spearman’s rank correlation *ρ* and its *P* value, calculated using 456 points, are shown in the upper-right corner. (**C**) Species–community interaction classes across 19 environments, indicated by point colour and plotted against normalised mono-culture cell density (x-axis) and normalised dropout-culture cell density (y-axis). Each cell-density value was normalised to the maximum for the same focal species or dropout-culture and displayed on a log_10_ scale. (**D**) Cumulative interaction-class counts across the same 152 combinations. Conditions are ordered from low to high raw endpoint cell density in mono-culture (upper plot) and dropout-culture (lower plot), with each step line representing one interaction class.

We next examined the association between endpoint cell density and interaction class. **Fig. 4C** shows interaction classes for individual conditions plotted against the normalised cell densities of the focal species in mono-culture and the resident community in dropout-culture. **Fig. 4D** shows cumulative interaction-class counts after ordering conditions by the corresponding raw endpoint cell density. To obtain a discrete summary, we divided the raw endpoint cell densities of the focal species in mono-culture and the resident community in dropout-culture at their respective medians, defining four cell-density quadrants (**Supplementary Fig. S6**). Overall, 94 of 152 interactions (61.8%) were classified as competition or amensalism. This combined competitive-interaction fraction was highest in the Focal High/Resident High quadrant, with 22 of 27 interactions (81.5%). A one-sided test showed an excess of competitive interactions in this quadrant relative to the overall fraction (environment-block permutation *P* = 0.04). The corresponding fractions were 9/27 (33.3%), 36/49 (73.5%), and 27/49 (55.1%) in the Focal Low/Resident Low, Focal High/Resident Low, and Focal Low/Resident High quadrants, respectively. Thus, competition and amensalism together were most frequent under conditions in which both the focal species and the resident community reached high endpoint cell densities.

### Multispecies communities buffer the growth enhancement associated with carbon-source mixing

In the previous study, mono-cultures tended to show greater growth in multi-carbon-source environments than the mean observed when the constituent carbon sources were supplied individually [24]. Having examined the association between endpoint growth and species–community interactions, we next tested whether this growth enhancement associated with carbon-source mixing persisted in multispecies communities. We compared the mean yield across three replicates in each multi-carbon-source environment with the mean of the corresponding single-carbon-source means, each based on three replicates (**Materials and Methods**). In mono-cultures, carbon-source mixing was associated with greater growth than expected from this mean, consistent with the previous study (**Fig. 5A**). This tendency was attenuated in multispecies communities, including dropout-cultures and the eight-species full-culture (**Fig. 5B**).

**Figure 5.**
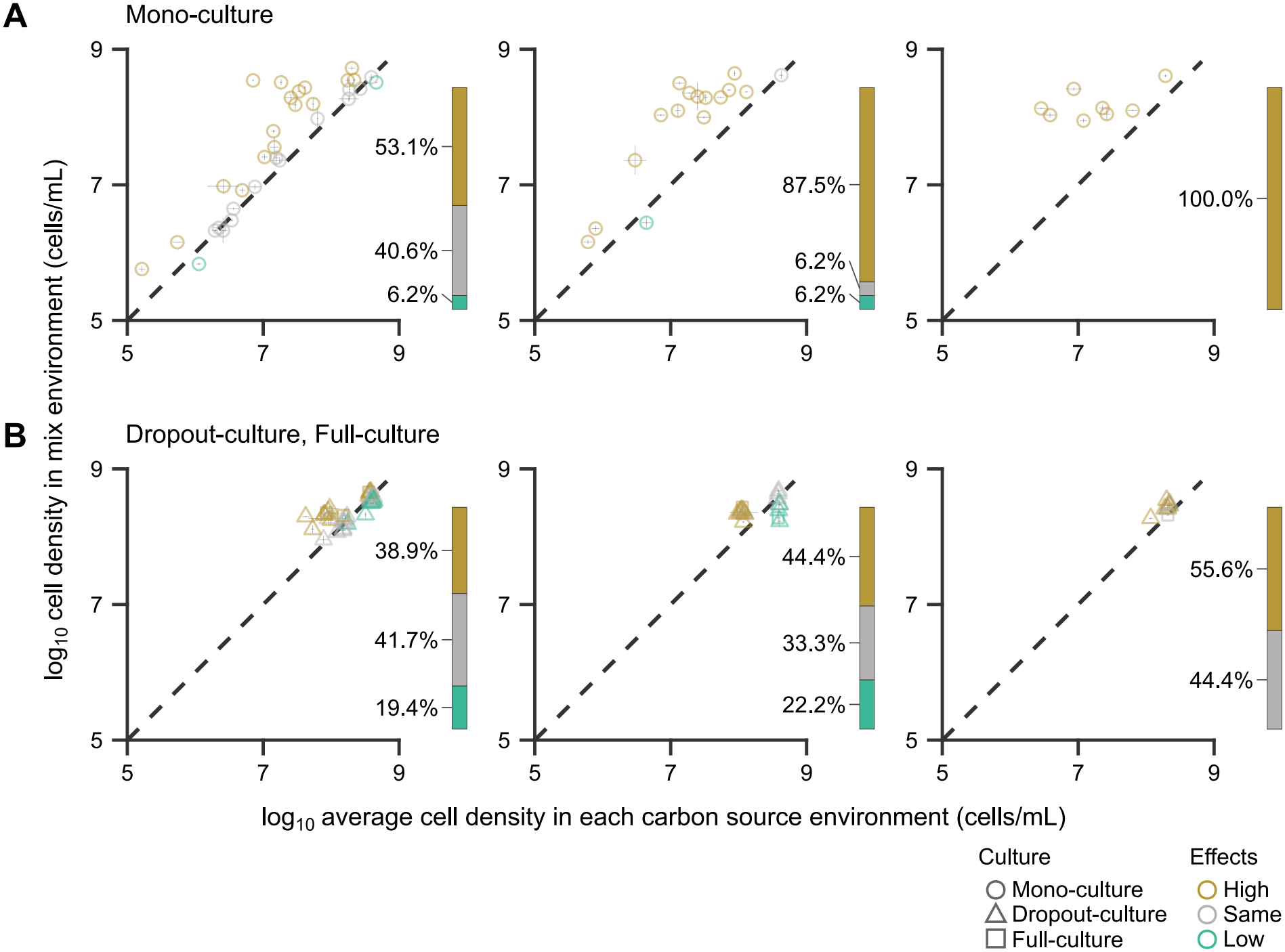
Growth enhancement associated with carbon-source mixing measured by cell density. Comparison of expected and observed cell densities following carbon-source mixing in mono-cultures (**A**) and dropout-cultures and full-cultures (**B**). Within each row, panels are arranged by target number of carbon sources (two, four, or eight). The x-axis shows the mean log_10_ cell density across the constituent single-carbon-source environments, and the y-axis shows the mean log_10_ cell density in the multi-carbon-source environment. Horizontal error bars show the square root of the mean of the within-environment sample variances, whereas vertical error bars show the sample standard deviation of the observed values in each multi-carbon-source environment. The dashed line indicates equality between expected (x-axis) and observed values (y-axis). Point shape indicates culture type, and colour indicates whether the observed cell density was higher than, the same as, or lower than expected from the single-carbon-source environments. The stacked sidebar beside each panel shows the fractions of High, Same, and Low among tested comparisons. The numbers of tested cases were 32 for two-carbon, 16 for four-carbon, and 8 for eight-carbon conditions in mono-culture and dropout-culture, and 4, 2, and 1, respectively, in full-culture. Four- and eight-carbon conditions were classified as Not tested when data were unavailable for one or more constituent single-carbon-source environments.

To quantify this attenuation directly, we first calculated the difference between log_10_ observed yield and log_10_ expected yield based on the constituent single-carbon-source environments, then averaged these differences within each culture group for each eligible target environment. We compared multispecies cultures with mono-cultures within the same environment using a paired t-test and a two-sided exact sign-flip test. Across seven eligible environments, growth enhancement was smaller in multispecies cultures than in mono-cultures (mean difference = −0.41 log_10_ units; 95% CI [−0.67, −0.15], paired-t *P* = 8.2 × 10^−3^, exact *P* = 0.02).

Carbon-source utilisation can affect cell size and morphology as well as cell number [56–58]. We therefore performed an analogous supplementary analysis using OD_595_ to capture such changes associated with carbon-source mixing. This analysis included 18 additional carbon-source environments beyond those used to investigate interspecies interactions (**Materials and Methods**; **Supplementary Table S2**). The OD_595_ analysis showed growth enhancement associated with carbon-source mixing in mono-cultures, with attenuation in dropout-cultures and the eight-species full-culture (**Supplementary Fig. S7**). The same direct contrast across 21 eligible target environments showed smaller growth enhancement in multispecies cultures than in mono-cultures (mean difference = −0.42 log_10_ OD_595_ units; 95% CI [−0.53, −0.31], paired-t *P* = 9.4 × 10^−8^, exact *P* = 2.9 × 10^−6^). Together, these results show that, although carbon-source diversification increases cell density and OD_595_ in mono-cultures, the associated growth enhancement is attenuated in multispecies communities.

Community yields in single-carbon-source environments were also compared with predictions from constituent mono-cultures. Observed yields were significantly higher than predicted in 60 of 72 community conditions (83.3%) for cell density and 65 of 72 conditions (90.3%) for OD_595_ (**Supplementary Fig. S8**).

### Species–species interactions show attenuated carbon-source dependence in multispecies communities

The species–community analysis showed that the carbon-source dependence of interspecies interactions observed in pairwise cocultures also occurs in multispecies communities. However, this definition does not reveal whether particular species drive the pattern or whether individual species pairs exhibit corresponding patterns. We therefore defined and analysed species–species interactions as net pairwise effect estimates for two species within the eight-species community. In growth environment *c*, the effect of target species *j* on focal species *i*, denoted by 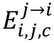, was quantified as the log_2_ ratio of full-culture to dropout-culture yield, using the mean yields of the focal species in the dropout-culture lacking the target species, 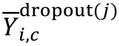, and full-culture, 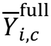 each based on three biological replicates. Conversely, the effect of focal species *i* on target species *j*, denoted by 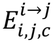, was quantified as the log_2_ ratio of full-culture to dropout-culture yield, using the mean yields of the target species in the dropout-culture lacking the focal species, 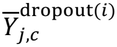, and full-culture, 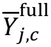, each based on three biological replicates (**Supplementary Fig. S9**). Like pairwise interactions measured in coculture, species–species interactions are bidirectional. The estimates are not directly equivalent, however, because species–species interactions include both direct effects between the focal and target species and indirect effects mediated by other species.

Species–species interactions were comparatively weak overall. Neutral interactions were the most frequent class, accounting for 172/224 (76.8%) pair–environment combinations in single-carbon-source environments (**Supplementary Fig. S10**) and 341/532 (64.1%) combinations across all 19 environments, including multi-carbon-source environments (**Supplementary Fig. S11**). Interaction strength, 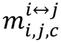, was calculated as the distance from the origin in the effect plane (**Materials and Methods**) and compared with matched prior pairwise data for the same environment and species pair. The current species–species interaction strength, 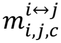, was smaller (mean difference = −1.75, crossed linear mixed-model 95% CI [−2.25, −1.26], likelihood-ratio test *P* = 5.6 × 10^−8^ ; **Supplementary Fig. S12A**). For descriptive reference, the distribution of current species–community interaction strengths is shown on the same y-axis scale (**Supplementary Fig. S12B**).

Although species–species interactions were weaker overall, they retained some structure. Despite the abundance of neutral interactions, the distribution of interaction classes varied with the number of carbon sources (**Supplementary Fig. S13A**). Interaction type, 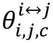, was negatively correlated with the number of carbon sources, although the correlation was weaker than those calculated for species–community interactions and pairwise cocultures (Spearman’s rank correlation: *ρ* = −0.17, *P* = 8.0 × 10^−5^, **Supplementary Fig. S13B**). A weak negative correlation was also observed after excluding neutral interactions (Spearman’s rank correlation: *ρ* = −0.28, *P* = 8.9 × 10^−5^, **Supplementary Fig. S13C**). In a supplementary analysis, we averaged interaction types across the 28 species-pair observations within each environment and re-evaluated the correlation using the 19 environment means. Neither the negative relationship based on all interactions in **Supplementary Fig. S13B** (Spearman’s rank correlation: *ρ* = −0.36, permutation *P* = 0.13; 95% environment-cluster bootstrap CI [−0.73, 0.15]) nor that after excluding neutral interactions in **Supplementary Fig. S13C** was significant (Spearman’s rank correlation: *ρ* = −0.23, permutation *P* = 0.34; 95% environment-cluster bootstrap CI [−0.62, 0.32]). We also compared these results with the previous study [24]; the prior–current correspondence table and direction-specific count audit are presented in **Supplementary Fig. S14** and **Supplementary Table S9**.

Together, these analyses show that species–species interaction strength in multispecies communities was lower than that of pairwise effects measured in pairwise cocultures, with many interactions operationally classified as neutral. Nevertheless, interactions did not disappear at random: some carbon-source and species-pair dependence remained in attenuated form.

## Discussion

In this study, we tested whether the shift towards competitive interactions observed in pairwise cocultures and the growth enhancement observed in mono-cultures with increasing carbon source diversity also occur in a synthetic bacterial community comprising eight species. As in the previous study [24], mono-cultures showed growth enhancement associated with carbon-source mixing, whereas this enhancement was attenuated in multispecies communities (**Fig. 5**). Species–community interactions nevertheless shifted towards competitive interactions as the number of carbon sources increased, consistent with the pattern observed in pairwise cocultures (**Fig. 3**). Thus, the shift towards competitive interactions with increasing carbon source diversity can persist even when growth enhancement associated with carbon-source mixing changes in multispecies communities.

In our previous study of pairwise cocultures, we suggested that two processes might contribute to the shift towards competitive interactions with increasing carbon source diversity: a greater likelihood of providing usable carbon sources to both species and growth enhancement associated with carbon-source mixing [24]. Here, the competitive shift persisted despite attenuated growth enhancement, suggesting that the relative importance of these processes may change with community complexity. Mono-culture growth of the focal species increased with carbon source diversity (**Fig. 4A**), whereas the total cell density of the resident community showed no corresponding increase (**Fig. 4B**). This difference may reflect the broad resource-utilisation capacity of the seven-species resident community, which allows some constituent species to grow even when few carbon sources are available. Increasing carbon source diversity may therefore increase the likelihood that the focal species can also grow sufficiently while the resident community maintains its growth. Consistent with this possibility, competitive interactions were more frequent when both groups showed high growth (**Fig. 4C and D**; **Supplementary Fig. S6**). Interaction patterns in complex communities may thus depend less on the specific metabolic processes underlying growth enhancement and more on statistical properties of resource availability, such as the likelihood that multiple community members can grow simultaneously. Paradoxically, increasing community complexity may thus simplify the drivers of competitive interactions, shifting explanatory power from species-specific metabolic responses to community-level statistical properties of resource availability.

The growth enhancement associated with carbon-source mixing observed in mono-cultures in the previous study [24] was attenuated in dropout-cultures and full-cultures (**Fig. 5**), showing that its extent differs between simple and complex species compositions. The mechanisms underlying this attenuation in multispecies communities remain unclear. Although experimental conditions, including total carbon-source concentration and the species examined, differ among studies, growth enhancement associated with combining carbon sources has been reported previously [24, 59, 60]. The specific mechanisms underlying this enhancement remain unresolved, but may include improved uptake or utilisation of supplied carbon sources through fulfilment of nutritional requirements [61, 62], relief of metabolic stress [63], and optimisation of metabolic networks [64]. Previous studies have linked microbial community composition to the diversity and composition of metabolites in the surrounding environment [28, 29]. In multispecies communities, the observed yield in single-carbon-source environments was already high relative to that of mono-cultures in the same environments (**Fig. 4A and B; Supplementary Fig. S8**), suggesting that constituent species collectively maintain sufficient growth even when carbon sources are limited. This may reflect the metabolic activities of different species generating diverse metabolites even when only one carbon source is supplied. These metabolites may support the growth of other community members, thereby substituting for the benefit of carbon-source mixing and reducing the additional growth enhancement observed in mono-cultures.

The species–community and species–species interactions evaluated here should be interpreted as net effects rather than measures of direct interactions alone. In species–community interactions, the effect of the resident community on the focal species integrates the net effects of the seven-species resident community. Conversely, the effect of the focal species on the resident community is its net effect on the total yield of that community. These estimates include indirect effects mediated by other species and are therefore not simple sums of direct interactions among individual species [25–27]. Similarly, species–species interactions include both direct effects between the focal and target species and indirect effects mediated by the other six species. The neutral category denotes the absence of a statistically significant effect, rather than the absence of an interaction.

This study shares several limitations with the previous study [24]. The environmental conditions were limited to M9-based medium, and the eight bacterial species examined were all Pseudomonadota that can be cultured without forming aggregates. Whether similar patterns occur under different nutrient conditions, pH-buffering capacities, temperatures, and phylogenetic contexts therefore warrants further study. Because interactions were evaluated after 72 hours of culture without serial passage, we could not assess temporal changes in interactions or long-term community dynamics. Nor can the data from this study alone directly identify the mechanisms underlying the observed interactions. The greater frequency of competitive interactions under conditions supporting high bacterial growth suggests that resource competition is a major molecular mechanism of interaction [16–19], but does not exclude other mechanisms. Metabolite exchange [20, 21] and secreted enzymes [65] may facilitate growth, whereas inhibitory-compound production [14, 15], as well as resource competition, may inhibit growth. The interactions evaluated here should therefore be interpreted as net effects arising from the simultaneous action of multiple mechanisms.

A limitation specific to this study is that species-specific absolute abundance was estimated from 16S rRNA gene amplicon sequencing data rather than measured directly. These estimates may be affected by species-specific biases in cell lysis, PCR amplification, and sequencing [66, 67]. In addition, because the total number of inoculated cells was held constant, the initial inoculum density per species differed among mono-culture, dropout-culture, and full-culture. The estimated net effects may therefore partly reflect differences in initial density. The strains also differed between studies: the previous study used fluorescently labelled strains, whereas the present study used the parental strains from which they were generated.

Despite these limitations, our results show that interactions can still shift towards competitive interactions with increasing carbon source diversity, even when growth enhancement associated with carbon-source mixing differs between mono-cultures and multispecies communities. The association between bacterial growth and competitive interactions identified in pairwise cocultures may therefore help explain some interaction patterns in multispecies bacterial communities. These findings provide a basis for relating environmental parameters to interaction patterns in complex microbial communities, suggesting that carbon source diversity acts as a robust driver of competitive interactions across scales of community complexity.

## Acknowledgments

We thank Tomoya Maeda (Hokkaido University and RIKEN, Japan) for providing the isolated environmental bacteria and Hazuki Kotani (RIKEN, Japan) for assistance with 16S rRNA gene amplicon sequencing analysis. Codex (GPT-6.1 Sol) was used to assist with code revision and improvement of the English writing.

## Author contributions

All authors contributed to the study design. H.O. conducted experiments, created visualisations, and wrote the original manuscript. H.O. and S.T. performed analyses. All authors reviewed and edited the manuscript. S.T. and C.F. supervised the project and acquired funding.

## Data availability

The datasets generated during this study are available on Zenodo [68]. The analysis codes are available on GitHub (https://github.com/HirokiOnoGit/Environment_and_multispecies_competition). The sequencing data have been deposited in the National Center for Biotechnology Information Sequence Read Archive (NCBI SRA) under BioProject accession PRJNA1560094.

## Supplementary material

supplementary-information20261112 - pdf file supplementary-tables20261112 - xlsx file

