## supplementary-information20261112 for "Carbon source diversity drives competitive shifts in multispecies bacterial communities despite attenuated growth enhancement"

### Title

Carbon source diversity drives competitive shifts in multispecies bacterial communities despite attenuated growth enhancement

### Short title

Environment and multispecies competition


### Authors

Hiroki Ono[1], Saburo Tsuru[2*], and Chikara Furusawa[2,3*]

### Affiliations

[1]Department of Biological Sciences, Graduate School of Science, The University of Tokyo, 7-3-1 Hongo, Bunkyo-ku, Tokyo 113-0033, Japan

[2]Universal Biology Institute, Graduate School of Science, The University of Tokyo, 7-3-1 Hongo, Bunkyo-ku, Tokyo 113-0033, Japan

[3]Center for Biosystems Dynamics Research, RIKEN, 6-7-1 Minatojima-minamimachi, Chuo-ku, Kobe 650-0047, Japan

### [*]Corresponding authors

Saburo Tsuru

Chikara Furusawa

**Contents**

**Supplementary Methods S1**. Reagent stock preparation

**Figure S1 Endpoint $OD_{595}$ stability**

**Figure S2 Raw abundance and community composition by condition**

**Figure S3 Fluorescence intensity gates and event distributions for total cell-density estimation**

**Figure S4 Relationship between $OD_{595}$ and cell density**

**Figure S5 Species–community interactions across carbon source diversity levels after excluding neutral interactions**

**Figure S6 Species–community interaction-class composition across endpoint cell-density quadrants**

**Figure S7 Growth enhancement associated with carbon-source mixing measured by $OD_{595}$**

**Figure S8 Community yields compared with mono-culture predictions in single-carbon-source environments**

**Figure S9 Schematic definition of species–species interactions**

**Figure S10 Species–species interactions in single-carbon-source environments**

**Figure S11 Species–species interactions across all pairs and environments**

**Figure S12 Weak interaction strengths in species–species interactions**

**Figure S13 Environmental structure in weak species–species interactions**

**Figure S14 Comparison of prior pairwise and current species–species interactions**

**Supplementary Methods S1**. Reagent stock preparation

Carbon-source stock solutions were prepared by dissolving the compounds in pure water at the following concentrations, then sterilised using a 0.20-μm filter and stored at −20 °C: D-glucose, 25.00 mg/mL; D-ribose, 25.00 mg/mL; D-cellobiose, 25.00 mg/mL; D-raffinose pentahydrate, 29.46 mg/mL; glycerol, 25.00 mg/mL; D-mannitol, 25.00 mg/mL; D-sorbitol, 25.00 mg/mL; sodium acetate, 34.73 mg/mL; trisodium citrate dihydrate, 38.88 mg/mL; disodium succinate hexahydrate, 58.19 mg/mL; L-alanine, 25.00 mg/mL; L-glutamine, 25.00 mg/mL; L-isoleucine, 25.00 mg/mL; L-proline, 25.00 mg/mL; L-serine, 25.00 mg/mL; and uridine, 25.00 mg/mL.

### Supplementary Figure Legends and Figures

#### Supplementary Figure S1. Endpoint $OD_{595}$ stability

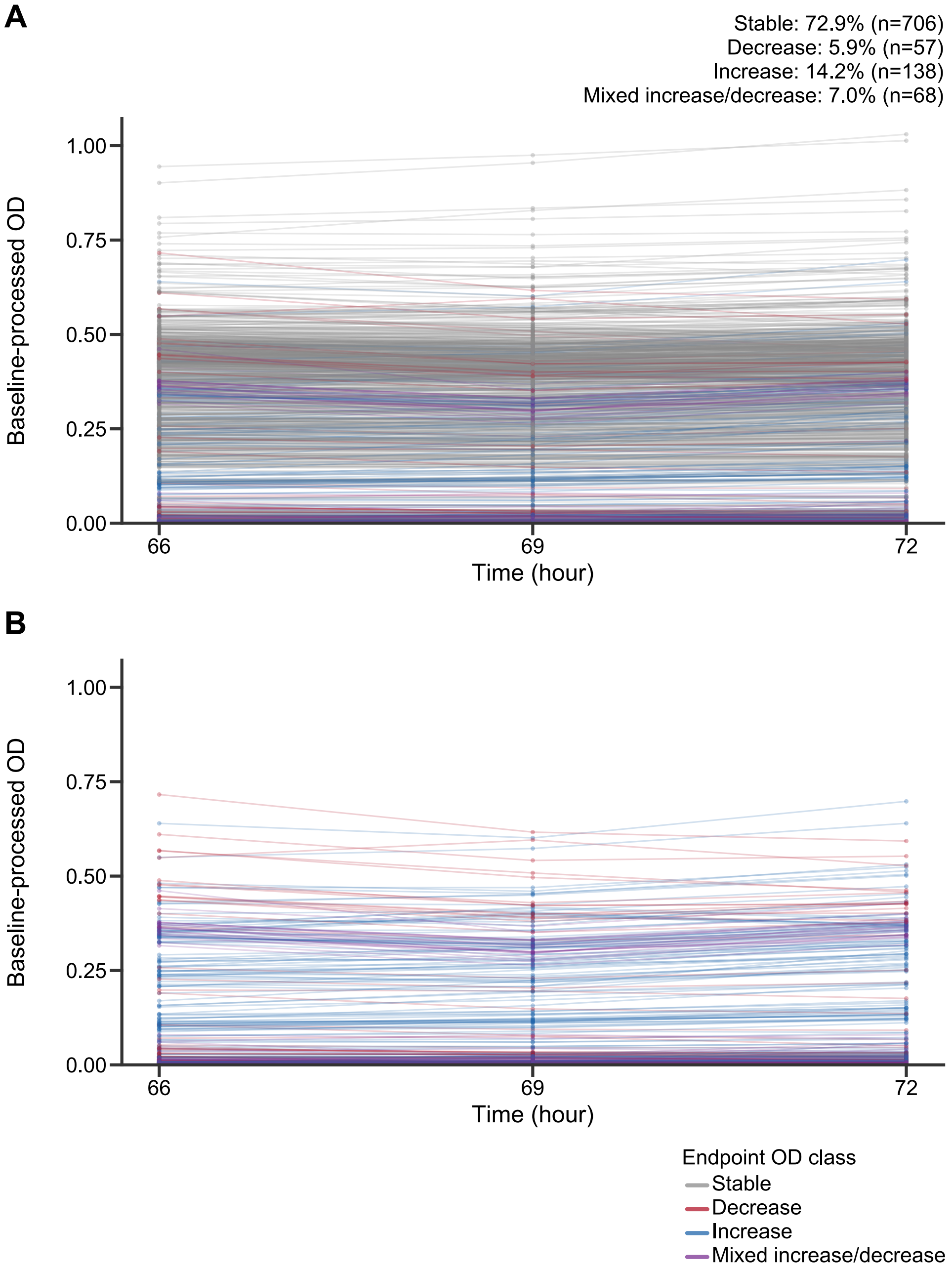


Changes in $OD_{595}$ at 66, 69, and 72 hours of culture, shown using blank-corrected values with measurements from the same well connected by lines. (**A**) All 969 samples from 19 environments × 17 compositions × three replicates. Line colour indicates whether $OD_{595}$ changed by more than 10% between 66 and 69 hours or between 69 and 72 hours, with percentage changes calculated relative to the higher $OD_{595}$ value in each interval. (**B**) Samples with a change of more than 10% in at least one interval.

### Supplementary Figure S2. Raw abundance and community composition by condition

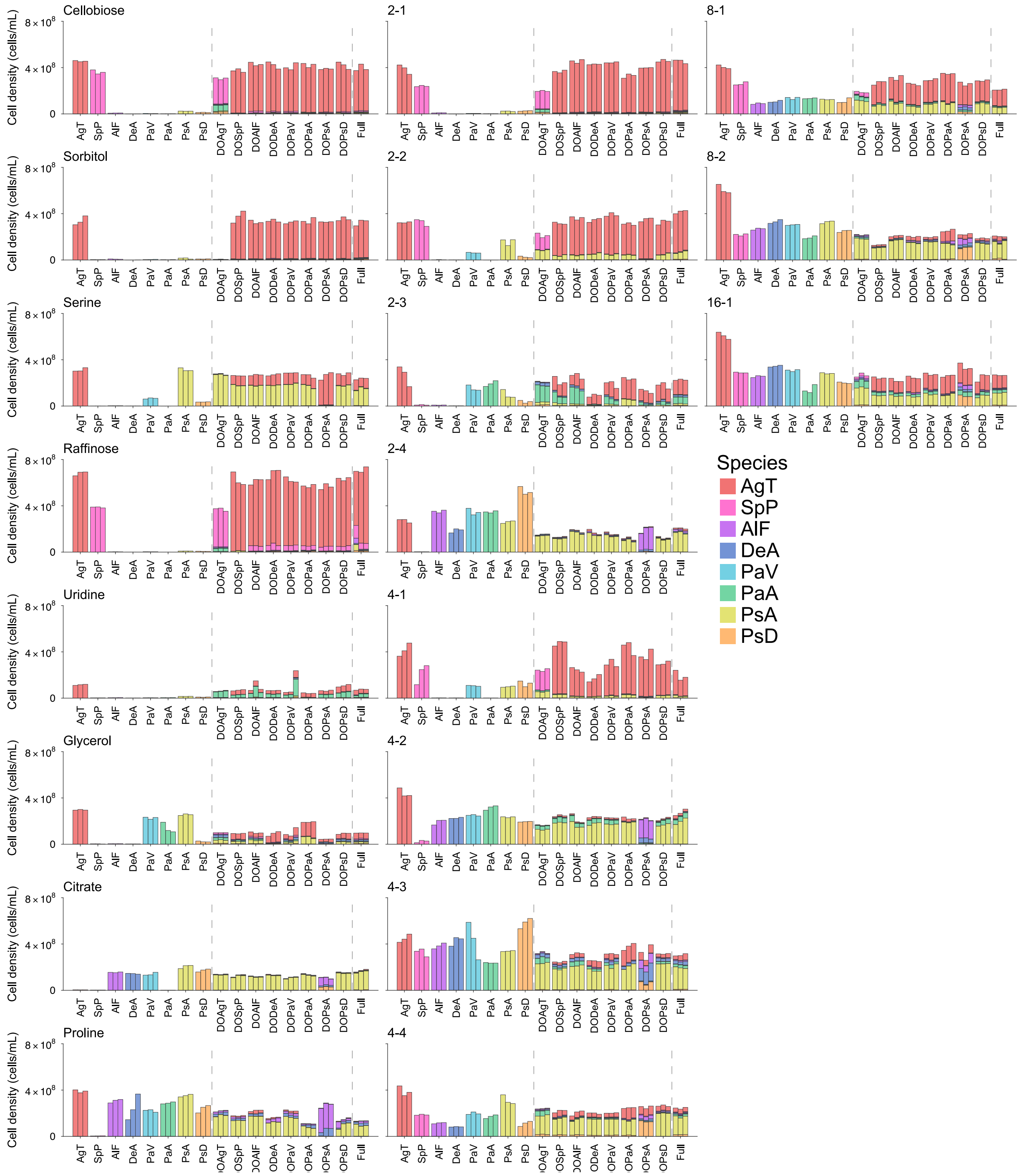


Species-specific absolute cell densities in each environment, with each subpanel representing one carbon-source environment. The eight bars on the left represent mono-cultures (AgT, SpP, AlF, DeA, PaV, PaA, PsA, and PsD); the central and right bars represent the dropout-cultures (DOAgT, DOSpP, DOAlF, DODeA, DOPaV, DOPaA, DOPsA, and DOPsD) and eight-species full-culture (Full), respectively. DO denotes dropout, and mono-culture bars show total cell density. For dropout-culture and full-culture, bar height is the sum of densities assigned to the displayed species, excluding the ambiguous group and omitted species without renormalisation; colours indicate species contributions.

Vertical dashed lines separate mono-culture, dropout-culture, and full-culture. Bacterial-species abbreviations are provided in **Fig. 1A**.

**Supplementary Figure S3. Fluorescence intensity gates and event distributions for total cell-density estimation**

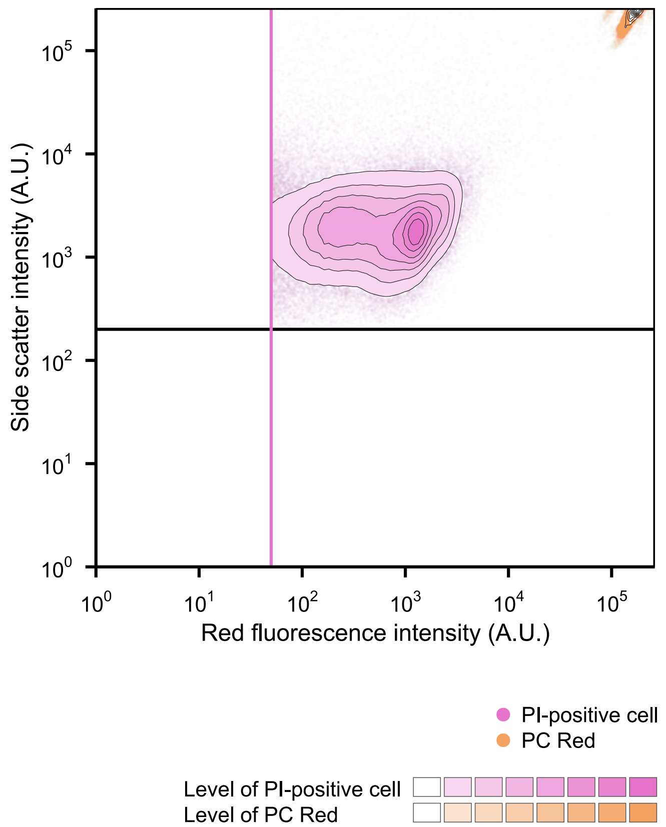


Overview of fluorescence intensity gates and event distributions. Events from all samples included in the analysis are overlaid on a plane defined by side scatter intensity and red fluorescence intensity, with pink indicating PI-positive cells and orange indicating PC Red. Events in each population were aggregated into two-dimensional bins and divided into eight density levels (Levels 1–8) on the basis of the maximum bin count for each population. Level 1 is transparent, with events selected for display in this region shown as scatter points. Levels 2–8 are filled with opaque colours, and black contours mark the density-level boundaries. The pink vertical line indicates the red fluorescence intensity threshold used to identify PI-positive cells, and the black horizontal line indicates the side scatter intensity threshold used to select events for display. Red fluorescence was excited with a 561-nm laser and measured in a detection band centred at 582 nm with a bandwidth of 15 nm (582 ± 7.5 nm). PC Red denotes Fluoresbrite® Polychromatic Red Microspheres (1.0 µm diameter).

**Supplementary Figure S4. Relationship between $OD_{595}$ and cell density**

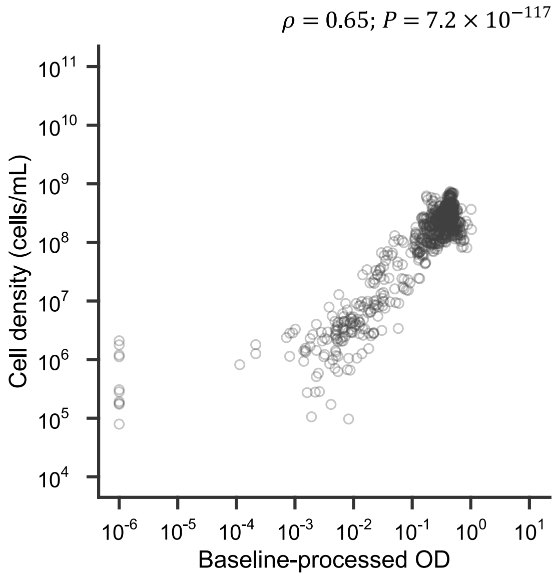


Association between $OD_{595}$ and cell density. Each point represents blank-corrected $OD_{595}$ and corrected cell density (cells/mL) for the same sample, with both axes displayed on a $\log_{10}$ scale. Spearman's rank correlation $\rho$ and its $P$ value are shown in the upper-right corner. Blank-corrected $OD_{595}$ values below $10^{-6}$ were set to this threshold. In a supplementary analysis, Spearman's rank correlation $\rho$ was calculated using 323 condition means, each based on three replicates, and evaluated using a block-permutation test in which cell-density condition means were permuted within each environment. This analysis also showed a positive relationship (Spearman's rank correlation: $\rho = 0.65$, block-permutation $P = 5.0 \times 10^{-5}$; 95% environment-cluster bootstrap CI $[0.38, 0.84]$).

**Supplementary Figure S5. Species–community interactions across carbon source diversity levels after excluding neutral interactions**

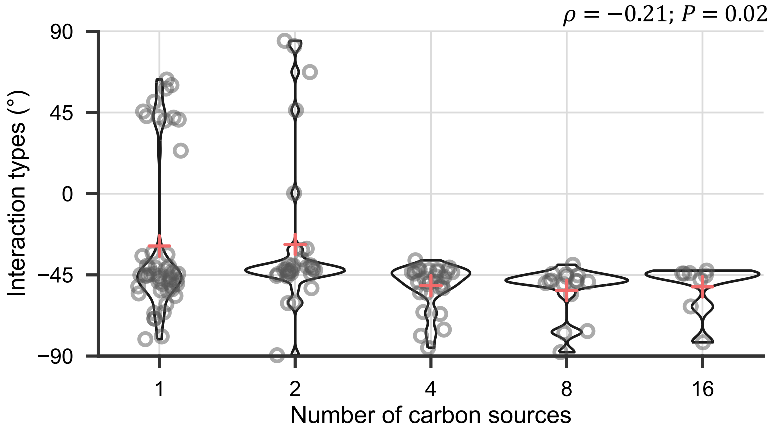


Jittered violin plots showing the distribution of interaction types, $\theta_{i,c}^{i\leftrightarrow\mathcal{R}(i)}$, for species–community interactions by number of carbon sources after excluding neutral interactions. Each point represents one focal species–environment combination, each violin shows the distribution, and cross-shaped points indicate the mean interaction type at each number of carbon sources. Spearman's rank correlation $\rho$ and its $P$ value, calculated using 133 points, are shown in the upper-right corner.

**Supplementary Figure S6. Species–community interaction-class composition across endpoint cell-density quadrants**

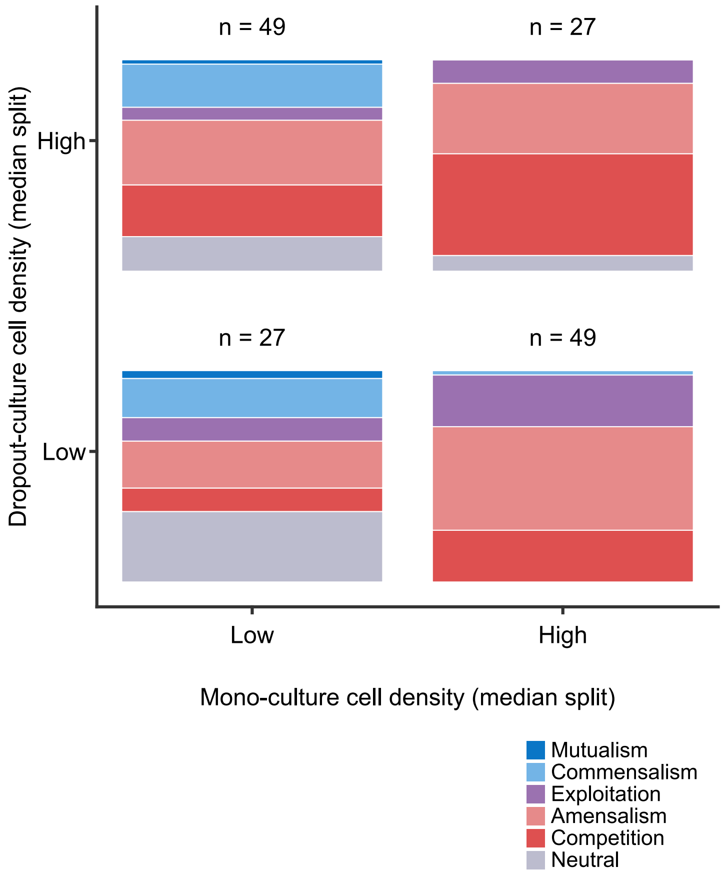


Interaction-class composition across 152 focal species–environment combinations in 19 environments. Raw endpoint cell densities at 72 hours of the focal species in mono-culture (x-axis) and the resident community in dropout-culture (y-axis) were each divided at their median across the 152 combinations. Each stacked tile shows the fractions of interaction classes within one quadrant, with the number of combinations indicated above.

**Supplementary Figure S7. Growth enhancement associated with carbon-source mixing measured by $OD_{595}$**

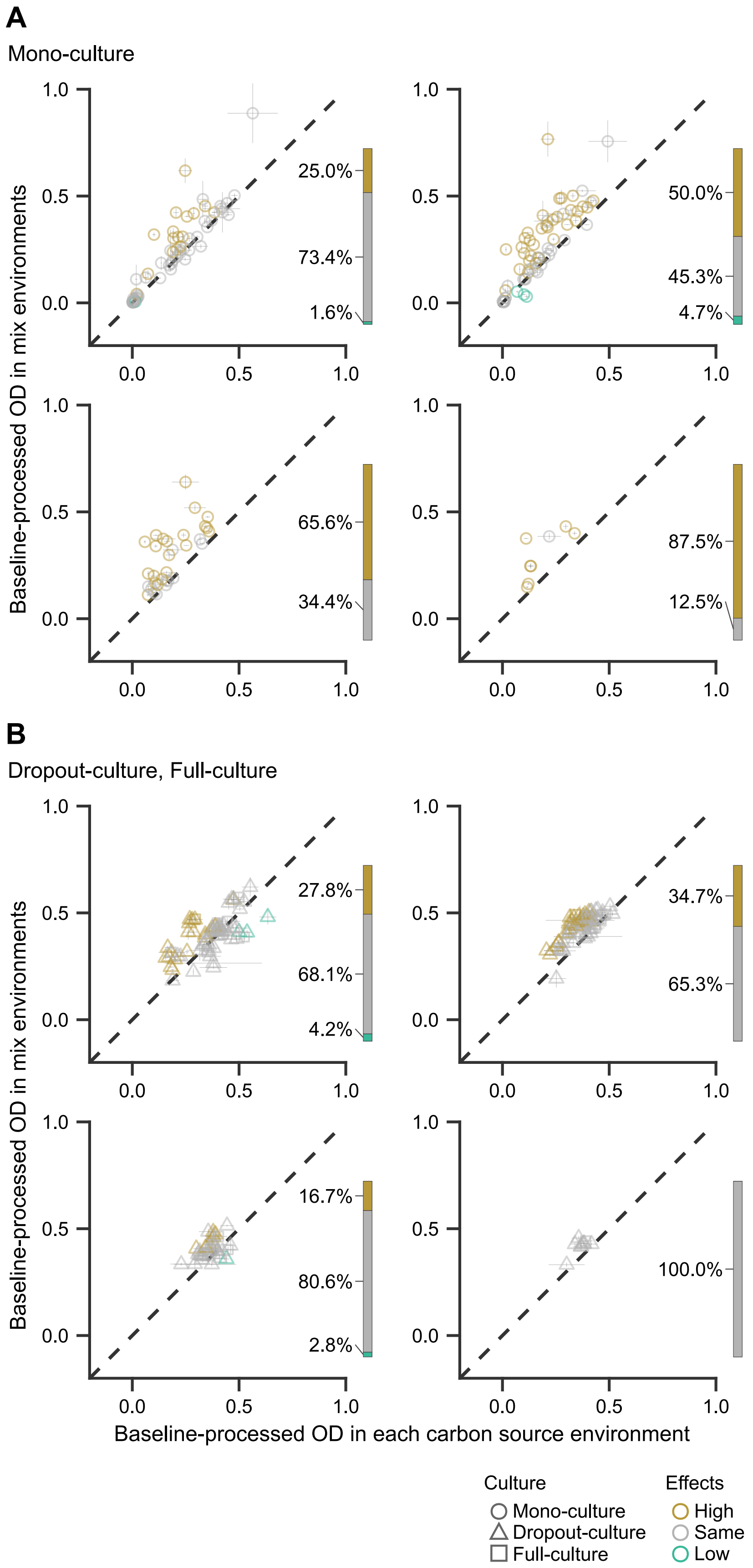

Comparison of expected and observed yields following carbon-source mixing in mono-cultures (**A**) and dropout-cultures and full-cultures (**B**) using $OD_{595}$ as the yield measure. Within each row, panels are arranged according to the target number of carbon sources: two (top left), four (top right), eight (bottom left), and 16 (bottom right). The x-axis shows the mean yield observed in the constituent single-carbon-source environments, and the y-axis shows the observed $OD_{595}$ in the multi-carbon-source environment. Horizontal error bars show the square root of the mean of the within-environment sample variances, whereas vertical error bars show the sample standard deviation of the observed values in each multi-carbon-source environment. The dashed line indicates equality between expected (x-axis) and observed values (y-axis). Point shape indicates culture type, and colour indicates whether the observed $OD_{595}$ was higher than, the same as, or lower than expected from the single-carbon-source environments. The stacked sidebar beside each panel shows the fractions of High, Same, and Low among the tests in that panel. The numbers of tested cases were 64 for two-carbon, 32 for four-carbon, 16 for eight-carbon, and 8 for 16-carbon conditions in mono-culture and dropout-culture, and 8, 4, 2, and 1, respectively, in full-culture.

**Supplementary Figure S8. Community yields compared with mono-culture predictions in single-carbon-source environments**

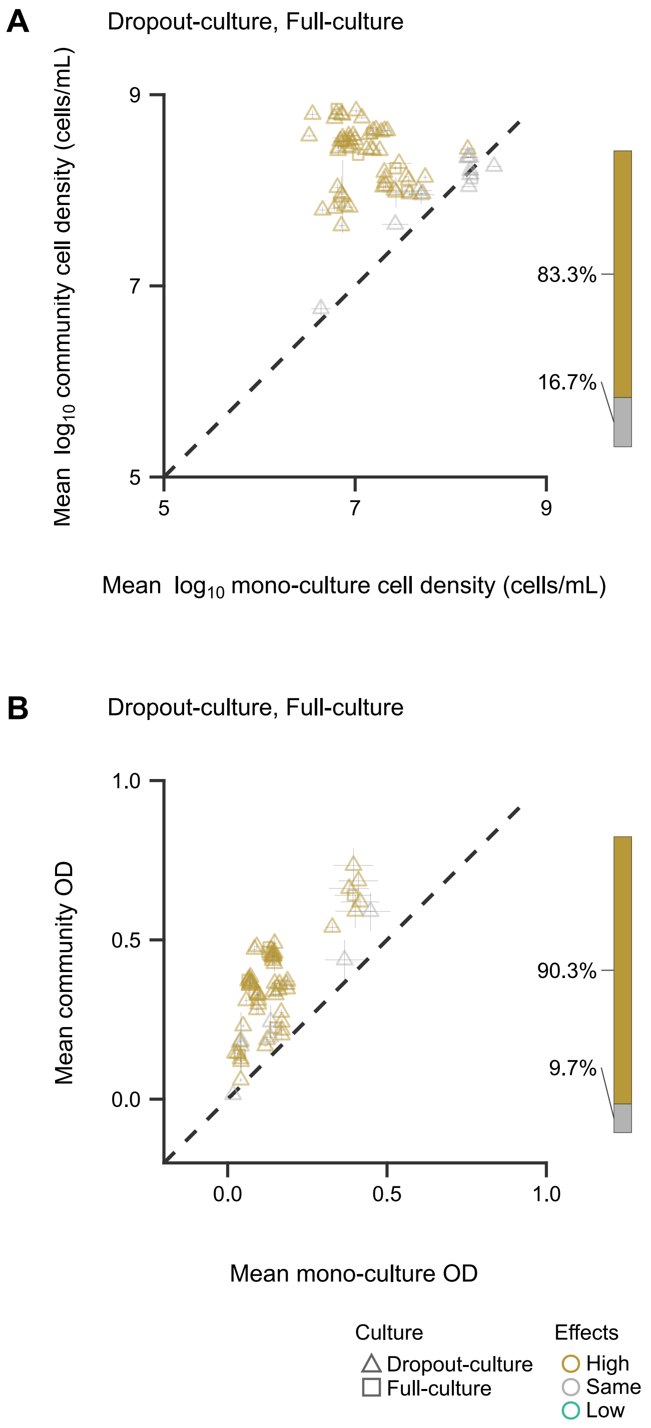


Comparison of predicted and observed community yields in eight single-carbon-source environments, measured by cell density on the $\log_{10}$ scale (**A**) and blank-corrected $OD_{595}$ on the original scale (**B**). Predictions are equal-weight means across the constituent species grown in mono-culture in the same environment. Triangles and squares indicate seven-species dropout-cultures and the eight-species full-culture, respectively. Each panel contains 72 community conditions (64 dropout-culture and eight full-culture conditions), each measured with three biological replicates. Horizontal error bars show the square root of the mean within-species sample variance; vertical error bars show the sample standard deviation of observed community replicates. Dashed lines indicate equality. Point colour indicates whether observed yields were significantly higher (High), lower (Low), or not significantly different (Same) from predictions (two-sided Welch *t*-tests against Group 2 defined in **Materials and Methods**). Stacked bars show the fractions of these classes across all 72 conditions.

### Supplementary Figure S9. Schematic definition of species–species interactions

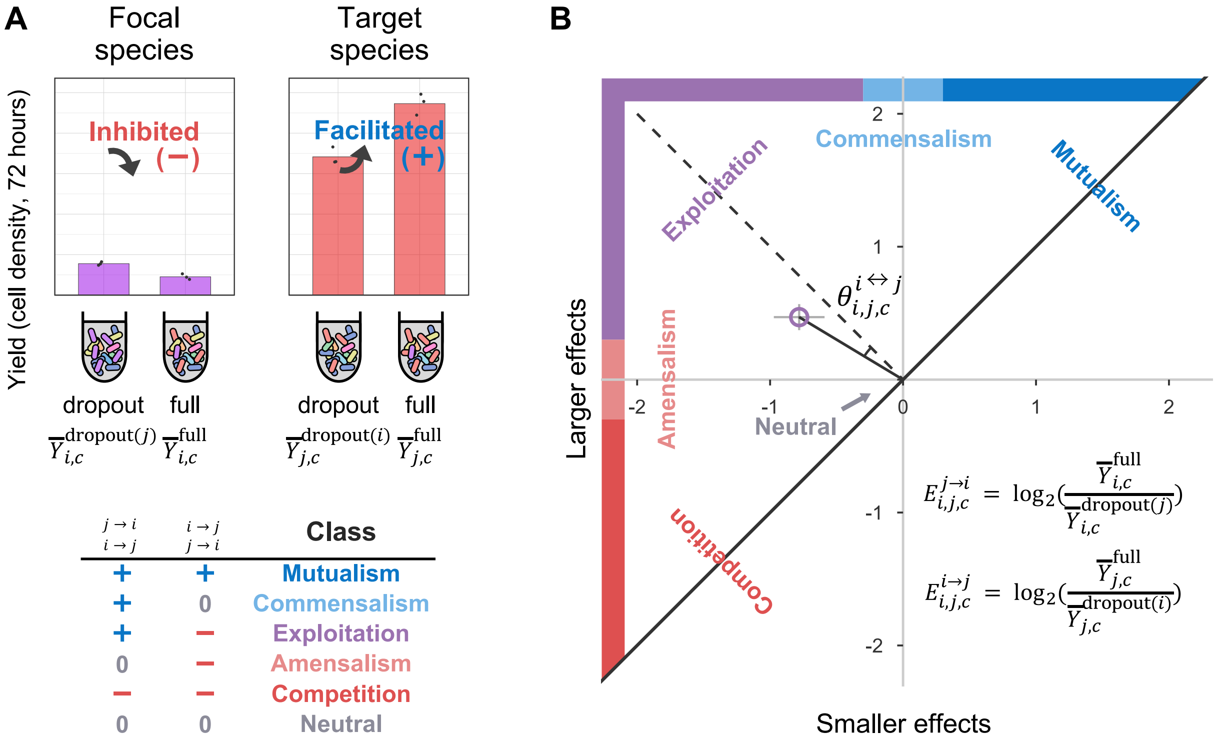


| $j \rightarrow i$<br>$i \rightarrow j$ | $i \rightarrow j$<br>$j \rightarrow i$ | Class |
|---|---|---|
| + | + | Mutualism |
| + | 0 | Commensalism |
| + | − | Exploitation |
| 0 | − | Amensalism |
| − | − | Competition |
| 0 | 0 | Neutral |

Conceptual framework for species–species interactions [1, 2]. (**A**, **B**) Evaluation of directional effects and bidirectional species–species interactions. For a dropout-culture lacking focal species $i$, the change in the yield of target species $j$ in full-culture relative to that dropout-culture defines the effect of the focal species on the target species, $E_{i,j,c}^{i\rightarrow j}$. Conversely, for a dropout-culture lacking target species $j$, the change in the yield of focal species $i$ in full-culture relative to that dropout-culture defines the effect of the target species on the focal species, $E_{i,j,c}^{j\rightarrow i}$. An interaction class was assigned from the signs of the two directional effects ($+, 0, -$; **Fig. 1C**). The larger and smaller of the two effects, namely the effect of the focal species on the target species, $E_{i,j,c}^{i\rightarrow j}$, and the effect of the target species on the focal species, $E_{i,j,c}^{j\rightarrow i}$, are plotted on the vertical and horizontal axes, respectively. The angle between the line segment connecting a plotted point to the origin and the line $y = -x$ defines interaction type, $\theta_{i,j,c}^{i\leftrightarrow j}$, whereas the distance from the origin defines interaction strength, $m_{i,j,c}^{i\leftrightarrow j}$. Error bars represent the standard deviation across three biological replicates.

**Supplementary Figure S10. Species–species interactions in single-carbon-source environments**

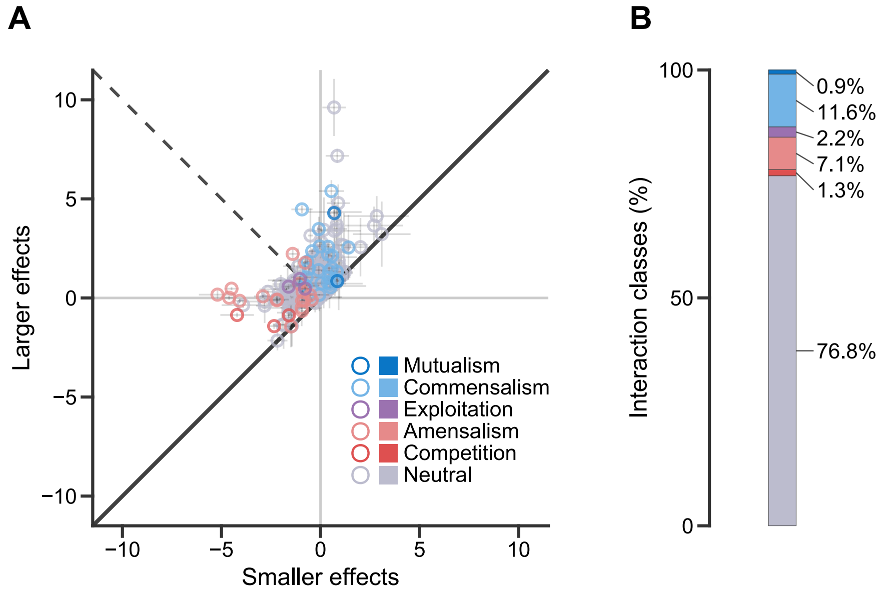


Species–species interaction patterns in single-carbon-source environments. (**A**) Effect plane in which each point represents one species-pair–environment combination. The x- and y-axes show the smaller and larger values of $E_{i,j,c}^{i\to j}$ and $E_{i,j,c}^{j\to i}$, respectively. Error bars indicate the standard deviation, and point colour indicates the assigned interaction class. (**B**) Fractions of interaction classes for the same single-carbon-source environments shown in **A**.

### Supplementary Figure S11. Species–species interactions across all pairs and environments

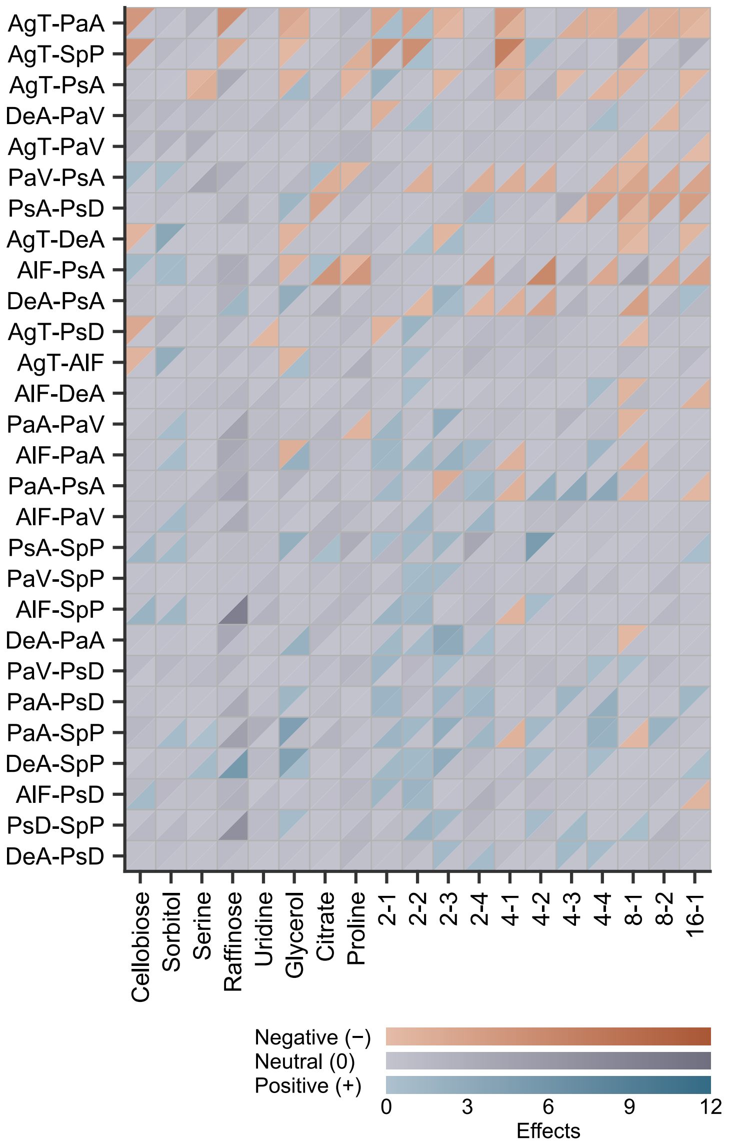


Split-triangle heatmap of species–species interactions for 28 species pairs across 19 environments, comprising 532 pair–environment combinations. The upper-left triangle of each cell indicates the effect of the species on the left of the pair label on the species on the right; the lower-right triangle indicates the reverse effect. Blue, red, and grey indicate positive, negative, and neutral effects, respectively; colour intensity indicates the magnitude of the corresponding directional $\log_2$ effect. Bacterial-species and carbon-source abbreviations are provided in **Fig. 1A** and **Table 1**, respectively.

**Supplementary Figure S12. Weak interaction strengths in species–species interactions**

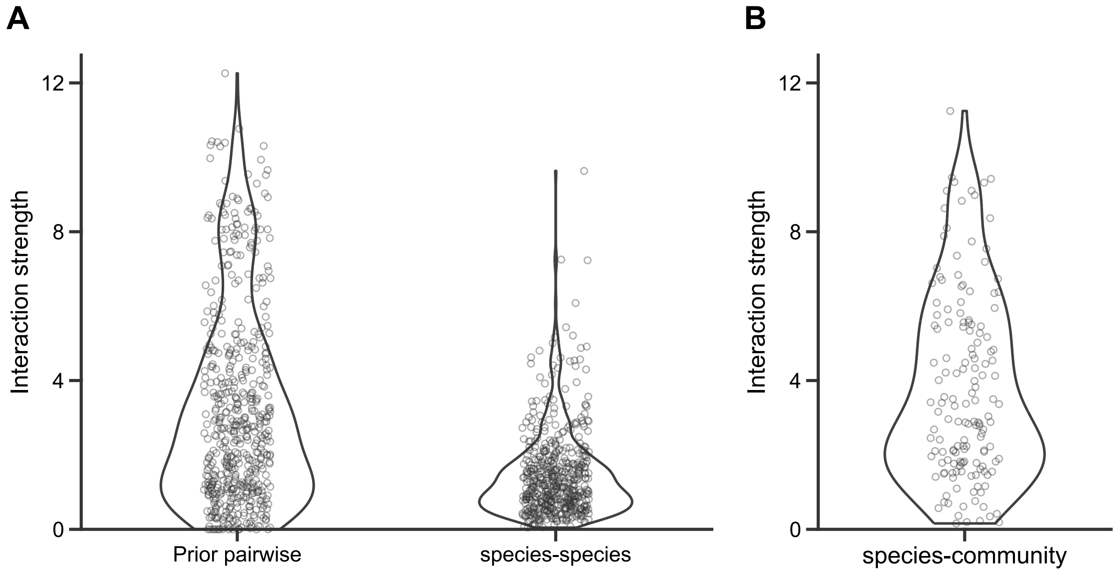


Comparison of interaction strengths among prior pairwise interactions [2], current species–species interactions, and current species–community interactions. Interaction strength was calculated as the Euclidean length of the vector comprising the two directional $\log_2$ effects. (**A**) Prior pairwise and current species–species interaction strengths for 532 environment × species-pair conditions (19 environments × 28 species pairs), with each point representing one condition. (**B**) Distribution of current species–community interaction strengths across 152 conditions, shown on the same y-axis scale as in **A**. Species–community interactions were not included in the test in **A**.

**Supplementary Figure S13. Environmental structure in weak species–species interactions**

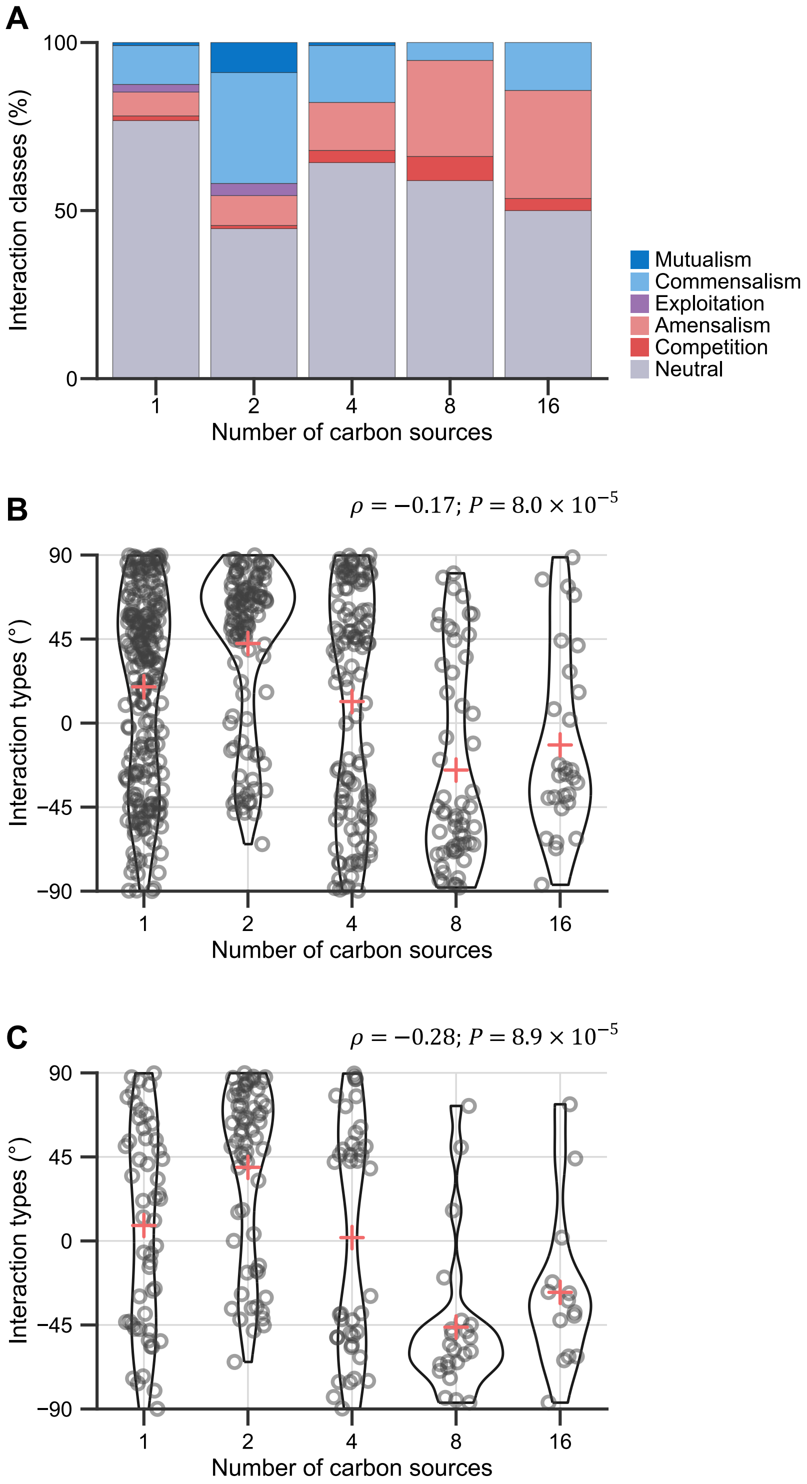


Environmental patterns in species–species interactions. (**A**) Interaction-class fractions across 19 environments, grouped by number of carbon sources. (**B**) Jittered violin plots showing the distribution of interaction types, $\theta_{i,j,c}^{i\leftrightarrow j}$, for the same species–species interactions by number of carbon sources. Each point represents one species-pair–environment combination, each violin shows the distribution, and cross-shaped points indicate the mean interaction type at each number of carbon sources. Spearman's rank correlation $\rho$ and its $P$ value, calculated using 532 points, are shown in the upper-right corner. (**C**) Jittered violin plots showing the distribution of interaction types, $\theta_{i,j,c}^{i\leftrightarrow j}$, after excluding neutral interactions from **B**. Spearman's rank correlation $\rho$ and its $P$ value, calculated using 191 points, are shown in the upper-right corner.

182 **Supplementary Figure S14. Comparison of prior pairwise and current species–species interactions**

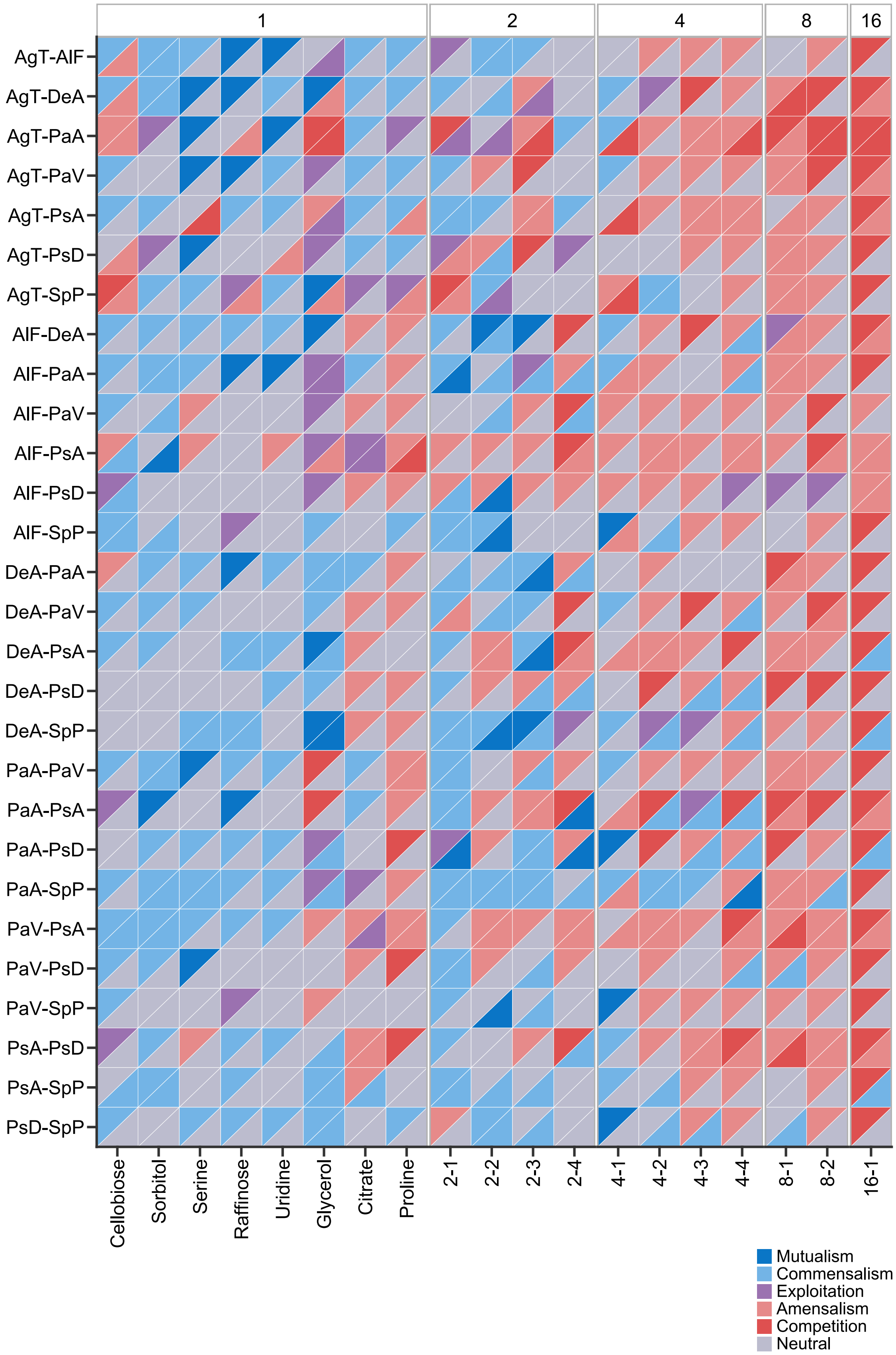


183

Concordance between prior pairwise and current species–species interaction classes [2], shown in a split-triangle heatmap. Each tile represents one species-pair–environment combination, with the upper-left triangle indicating the prior pairwise interaction and the lower-right triangle indicating the current species–species interaction. Columns represent environments grouped by number of carbon sources (one, two, four, eight, or 16), and rows represent species pairs. Colour indicates interaction class. Bacterial-species and carbon-source abbreviations are provided in **Fig. 1A** and **Table 1**, respectively.
